# The inflammatory microenvironment of the lung at the time of infection governs innate control of SARS-CoV-2 replication

**DOI:** 10.1101/2024.03.27.586885

**Authors:** Paul J. Baker, Andrea C. Bohrer, Ehydel Castro, Eduardo P. Amaral, Maryonne Snow-Smith, Flor Torres-Juárez, Sydnee T. Gould, Artur T. L. Queiroz, Eduardo R. Fukutani, Cassandra M. Jordan, Jaspal S. Khillan, Kyoungin Cho, Daniel L. Barber, Bruno B. Andrade, Reed F. Johnson, Kerry L. Hilligan, Katrin D. Mayer-Barber

## Abstract

SARS-CoV-2 infection leads to vastly divergent clinical outcomes ranging from asymptomatic infection to fatal disease. Co-morbidities, sex, age, host genetics and vaccine status are known to affect disease severity. Yet, how the inflammatory milieu of the lung at the time of SARS-CoV-2 exposure impacts the control of viral replication remains poorly understood. We demonstrate here that immune events in the mouse lung closely preceding SARS-CoV-2 infection significantly impact viral control and we identify key innate immune pathways required to limit viral replication. A diverse set of pulmonary inflammatory stimuli, including resolved antecedent respiratory infections with *S. aureus* or influenza, ongoing pulmonary *M. tuberculosis* infection, ovalbumin/alum-induced asthma or airway administration of defined TLR ligands and recombinant cytokines, all establish an antiviral state in the lung that restricts SARS-CoV-2 replication upon infection. In addition to antiviral type I interferons, the broadly inducible inflammatory cytokines TNFα and IL-1 precondition the lung for enhanced viral control. Collectively, our work shows that SARS-CoV-2 may benefit from an immunologically quiescent lung microenvironment and suggests that heterogeneity in pulmonary inflammation that precedes or accompanies SARS-CoV-2 exposure may be a significant factor contributing to the population-wide variability in COVID-19 disease outcomes.

## INTRODUCTION

By the end of 2021, half of the global population had been infected at least once with SARS-CoV-2 (hereafter SCV2) ^1^, the causative virus of COVID-19, with striking population-wide variability in disease outcome. Prognoses include asymptomatic infection, mild, non-specific pulmonary symptoms with or without the development of post-acute sequelae of COVID-19 or severe respiratory distress requiring hospitalization and mechanical ventilation, which can lead to death. Better understanding of the genetic and immunological determinants of heterogenous disease outcomes could lead to measures that improve the clinical management of COVID-19.

Contributing factors to the diversity in clinical outcomes include infectious dose and viral strain differences, alongside host factors like age, sex, genetics, and vaccination status, as well as defects in innate antiviral immunity or underlying comorbidities including obesity and diabetes ^2–16^. Human genetic variation associated with disease severity has revealed defective innate immune responses as another key determinant in COVID-19 disease outcomes. Patients with inborn errors in components of RNA-sensing and innate signaling pathways, including toll-like receptor 3 (TLR3), TLR7, interferon-regulatory factor 7 (IRF7), type-I IFN (IFN-I) receptors IFNAR1 and IFNAR2, tyrosine kinase 2 (TYK2) and oligoadenylate synthetase 1 (OAS1) are at high risk of developing critical and severe COVID-19 disease ^2–4,15,17–19,12,20–22^. The crucial importance of innate-derived IFNs is further underscored by the discovery that up to 15 – 20% of critically ill patients with COVID-19 had preexisting auto-antibodies to IFN-I, which delayed viral clearance ^23–26^. Thus, during the early phase of SCV2 infection, defective innate immune responses due to genetic defects or auto-autoantibodies fundamentally influence disease outcome ^3,27^. Understanding pulmonary innate antiviral responsiveness prior to SCV2 exposure may, therefore, shed further light on the variability in clinical presentations.

Patient populations with pre-existing pulmonary diseases, although initially predicted to be more vulnerable to SCV2 infection, have unexpected heterogeneity in COVID-19 outcomes. While certain chronic lung diseases, including tuberculosis (TB) ^28,29^ and chronic obstructive pulmonary disease (COPD) ^30–33^ have been associated with increased severity of COVID-19 in most studies, other chronic pulmonary conditions, such as asthma ^32–35^ and cystic fibrosis (CF) ^36–38^ did not consistently correlate with worsened COVID-19 presentation and have even been associated with improved disease outcomes. The immunological factors in the lung that determine such variability in early viral control, and thus the likelihood of developing severe disease, are incompletely understood, and challenging to examine in clinical settings.

Here, we demonstrate using experimental mouse models of respiratory SCV2 infection that recent pulmonary bacterial or viral infections or underlying allergic inflammation precondition the lung for enhanced control of SCV2 replication. Importantly, administration of individual TLR9 or TLR1/2 ligands, recombinant TNFα or recombinant IL-1 to the lung prior to SCV2 infection also resulted in lower viral titers early after infection. Additionally, we surveyed pulmonary innate inflammatory pathways and identified a range of innate immune signaling components necessary for controlling early SCV2 replication in the lungs of mice. Collectively, our work reveals that the inflammatory lung microenvironment during SCV2 exposure may be a previously underappreciated, important factor in influencing disease variability and outcome through potent innate restriction of early viral replication.

## RESULTS

### Recent infection or underlying inflammation of the lung at the time of SARS-CoV-2 exposure limits pulmonary viral replication

To explore whether the recent infectious and inflammatory history of the lung could impact early viral replication, we exposed mice to a variety of respiratory pathogens or sterile inflammatory stimuli prior to SCV2 infection. For example, we and others have previously demonstrated that ongoing presence of mycobacteria in the lungs after infection with chronic mycobacterial pathogens such as *Mycobacterium tuberculosis* (*Mtb)* or Bacille Calmette-Guérin (BCG) results in lower viral titers and protection against SCV2 ^39–44^. C57BL/6 wild type (WT) mice were infected with *Mtb* 3 – 4 months prior to infection with SCV2 variant of concern (VOC) B.1.351 (beta variant) **(Fig 1A)**. Consistent with our previous data, mice with an ongoing *Mtb* infection exhibited significantly decreased lung viral titers three days post SCV2 infection compared to mice without underlying infection as detected by both TCID_50_ assay and qPCR for the SCV2 envelope (E) gene in its actively replicating form, (subgenomic, sub-gRNA) **(Fig 1A)**. To investigate whether suppression of SCV2 replication is specific to mycobacterial coinfections, we next exposed mice to Methicillin-resistant *Staphylococcus aureus* (*S. aureus*, USA300), a gram-positive bacterial pathogen that is a major cause of nosocomial infections ^45^. While intratracheal inoculation *of S. aureus* induces a potent immune infiltrate in the lungs, it is rapidly cleared within 48 hours in mice ^46^. Taking advantage of the rapid bacterial clearance in this model, we tested whether recent inflammation elicited in response to extracellular bacteria is equally protective as actively replicating intracellular mycobacteria. Indeed, when we intrapharyngeally (i.ph.) exposed the lungs of WT mice to *S. aureus* three days prior to SCV2 B.1.351 infection, mice with recently cleared *S. aureus* infection exhibited significantly lower SCV2 viral titers than mice without prior exposure to *S. aureus* **(Fig 1B)**. Thus, both very recent and chronic pulmonary bacterial infections can promote an antiviral state in the mouse lung that lowers SCV2 titers prior to the onset of adaptive immunity and this protective feature is not unique to mycobacteria.

**Figure 1:**
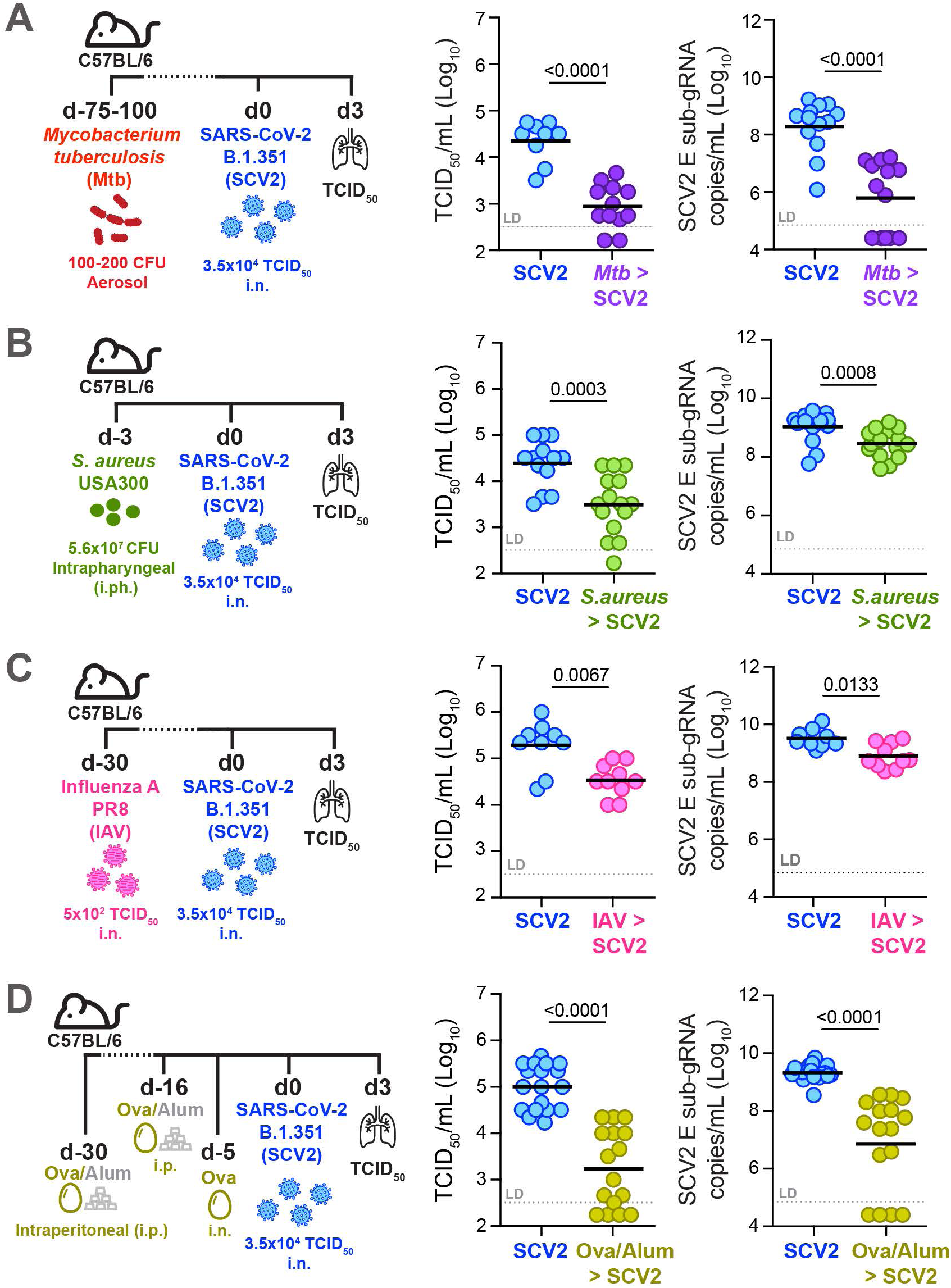
Recent respiratory infection or underlying pulmonary inflammation at the time of SCV2 exposure limits early viral replication in the lungs of mice. For SCV2 (SCV2) infections, all mice were infected intranasally (i.n.) with 3.5×10^4^ TCID_50_ SCV2 (B.1.351) and euthanized three days later (d3). Viral loads were measured by TCID_50_ on Vero E6 cells and by qPCR for the SCV2 E gene in its sub-genomic form (sub-gRNA) **(A)** Lung viral loads of WT mice infected with *Mtb* by aerosol exposure 2 – 4 months before infection with SCV2 **(B)** Lung viral loads of WT mice infected intraphayngeally (i.ph.) with *S. aureus* USA300 three days before SCV2 infection **(C)** Lung SCV2 loads of WT mice i.n. infected with Influenza A virus (IAV, PR8) one month before SCV2 infection **(D)** Lung viral loads of WT mice intraperitoneally (i.p.) injected twice with ovalbumin and aluminum hydroxide (ova-alum) 30 and 16 days before SCV2 infection and i.n. OVAwas given 5 days before SCV2 infection. n= 9 – 18, data combined from 2 – 3 independent experiments, geometric mean, statistical significance calculated by two-tailed Mann Whitney test, LD= limit of detection.

We then asked whether this innate antiviral state is specific to previous bacterial lung infections or whether prior lower-respiratory viral infections can similarly confer improved innate viral control in the lungs of mice. Simultaneous co-infection with Influenza A Virus (IAV) has been shown in both *in vitro* and *in vivo* studies to potently interfere with SCV2 replication, a phenomenon known as ‘viral interference’ ^47–51^, but it remains unclear whether a recently resolved and cleared IAV infection could also affect early viral SCV2 replication. Thus, rather than simultaneously co-infecting mice, we intranasally (i.n.) infected with IAV one month prior to SCV2 infection, a time frame by which IAV has been cleared for a minimum of two weeks ^52,53^. Mice that had recently cleared IAV infection displayed a significant reduction in lung SCV2 viral titers compared to mice without a recent IAV infection **(Fig 1C)**. These data suggest that the pulmonary anti-SCV2 state induced by prior pathogen exposure does not require ongoing infection and can persist for at least two weeks after prior pathogen clearance.

Finally, to determine whether only prior live pathogens and/or type I immune responses are necessary to restrict SCV2 replication in the mouse lung, we induced a sterile type II immune-driven allergic inflammatory response using the ovalbumin (OVA)/alum-driven asthma model. Five days after i.n. challenge with OVA, mice were infected with SCV2 B.1.351 and we found that mice with underlying type II inflammation displayed a significantly reduced lung burden of SCV2 **(Fig 1D)**. These results establish that both live pathogen and sterile type I- or type II-driven, recent, or ongoing inflammatory responses restrain initial SCV2 replication in the mouse lung and suggest that the recent pulmonary exposure history contributes to disease trajectory.

### Previous pulmonary TLR stimulation is sufficient to suppress SCV2 replication

While we showed that recent diverse pulmonary exposures, ranging from pathogens to sterile asthmatic inflammation, condition the lung for improved viral SCV2 control, the precise underlying cellular and molecular mechanisms in each setting are likely very complex. Therefore, we evaluated whether a single pulmonary administration of a TLR ligand, one week prior to SCV2 infection would be sufficient to promote an antiviral state in the lungs of mice, allowing for more amenable exploration of potential mechanisms that contribute to viral restriction. Such an approach would also inform on the effects of TLR agonists used as adjuvants in conjunction with antigens in mucosally delivered vaccines. Importantly, we found that i.ph. administration of the TLR9 agonist type B CpG (CpG) or the TLR1/TLR2 agonist Pam3CSK4 (Pm3) to WT mice one week prior to SCV2 B.1.351 infection, resulted in a significant reduction in SCV2 burden compared to mice that were not pre-treated with TLR ligands **(Fig 2A**, **S1A)**. To investigate the duration of protection afforded by one-time prior TLR activation in the lung following CpG administration, rather than seven days, we rested mice for seven weeks before infecting them with SCV2. Remarkably, the protection afforded by prior pulmonary CpG administration did not persist, as lung SCV2 titers were unchanged between CpG pre-treated and untreated mice seven weeks after CpG preconditioning **(Fig 2B**, **S1B)**. Thus, only recent pulmonary exposure to the TLR9 agonist CpG provided early viral replication control.

**Figure 2:**
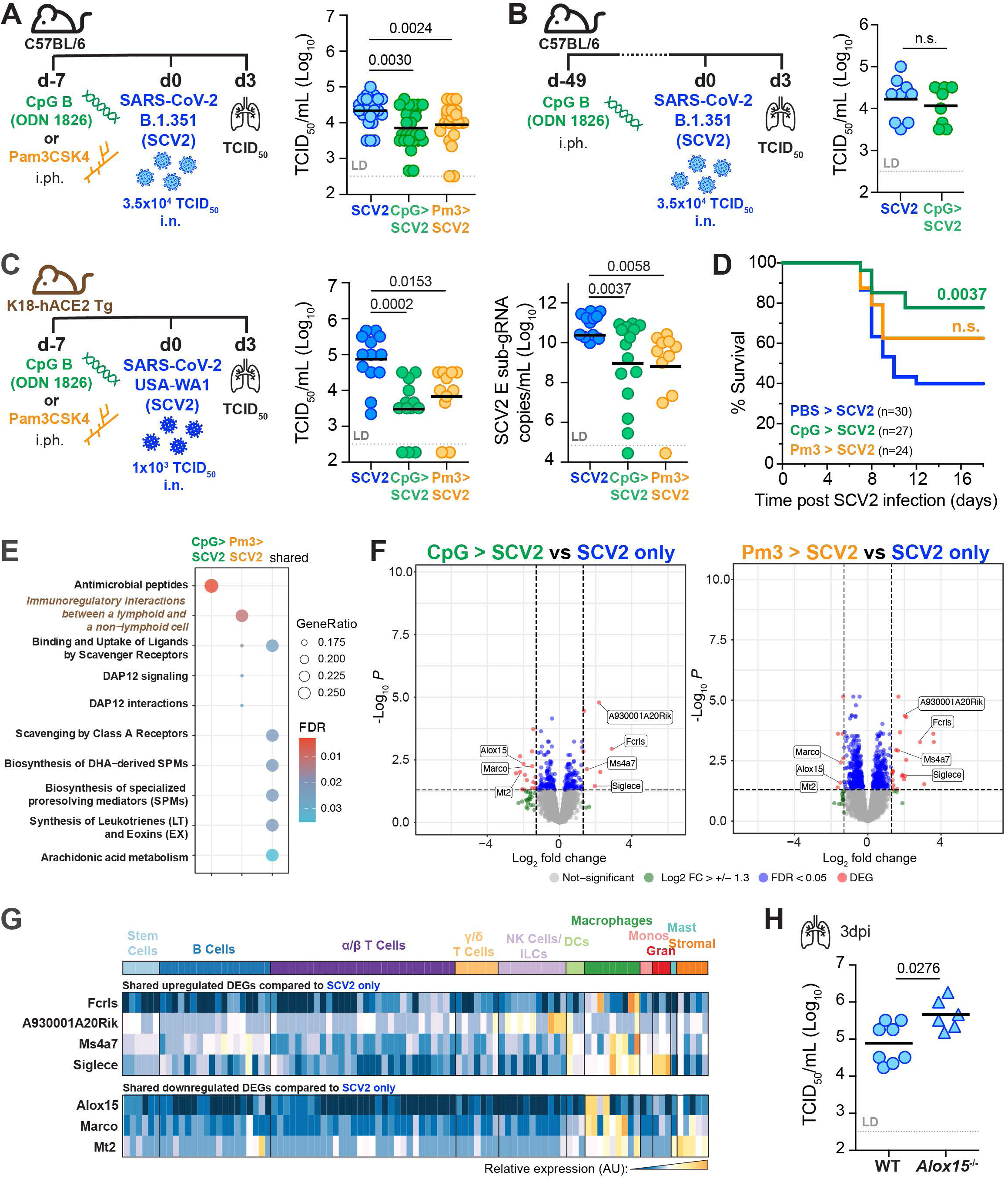
Recent one-time pulmonary TLR pre-stimulation is sufficient to suppress early SCV2 replication in the lung. **(A)** Left: WT mice were administered PBS, 10μg CpG B (ODN1826, CpG) or 50μg Pam3CSK4 (Pm3) i.ph. one week before i.n. infection with 3.5×10^4^ TCID_50_ SCV2 (SCV2) (B.1.351) and assessed for lung viral loads three days later (3dpi). Right: TCID_50_ lung viral loads n=20-26, data combined from five independent experiments **(B)** Left: WT mice were i.ph. administered PBS or 10μg CpG i.ph seven weeks before i.n. infection with 3.5×10^4^ TCID_50_ SCV2 (B.1.351). Right: TCID_50_ lung viral loads at 3dpi, n= 8-9, data combined from two independent experiments. **(C)** Left: K18-hACE2 Tg mice were i.ph. administered PBS, 10μg CpG or 50μg Pm3 one week before i.n. infection with 1×10^3^ TCID_50_ SCV2 (USA-WA1/2020) Right: lung viral loads 3dpi measured by TCID_50_ or qPCR for sub-gRNA SCV2 E gene, n= 11-13, data combined from three independent experiments. **(D)** K18-hACE2 Tg mice were treated and infected as described in (C) and monitored for time to clinical endpoint (survival) for 18 days post SCV2 infection. Mouse survival is shown as a Kaplan-Meier curve with significance determined by Mantel-Cox test, n= 24-33, data combined from six independent experiments **(E – G).** K18-hACE2 Tg mice i.ph. administered PBS, CpG or Pm3 and one week later infected i.n. with SCV2 USA-WA1/2020 Lung total RNA sequencing was performed at 3dpi (n = 3-4 mice per group in one experiment). **(E)** GO analysis of significant DEGs **(F)** Volcano plots of candidate DEGs comparing SCV2 infected mice pre-treated with CpG (left panel) or Pm3 (right panel) to SCV2-only infected control animals. DEGs significantly upregulated in both treatment groups are labeled. **(G)** DEGs after SCV2 infection in common to both TLR pretreatment groups were entered into ImmGen MyGeneSet. Expression across cell types as analyzed by ImmGen are visualized in a heatmap, AU (arbitrary units), navy= lowest expression, orange= highest expression; ILCs (innate lymphoid cells), DCs (dendritic cells), Monos (monocytes), Grans (granulocytes), Mast (mast cells) **(H)** Viral loads in lungs of WT and *Alox15*^−/−^ mice 3 days post-i.n. infection with 3.5×10^4^ TCID_50_ SCV2 (B.1.351) as measured by TCID_50_ on Vero E6 cells, n= 6-8, data combined from 2 independent experiments. **(A – C, H)** geometric mean, statistical significance determined by two-tailed Mann Whitney test, LD= limit of detection, n.s.= not significant.

To extend our observations to another mouse model of SCV2 infection and to rule out the possibility that our findings may be unique to infections of C57BL/6 mice with the SCV2 VOC B.1.351, we similarly pre-exposed lungs of mice transgenically expressing human ACE2 under the control of the epithelial K18 promoter (K18-hACE2 Tg) mice to a single dose of CpG or Pm3 one week prior to infection with the ancestral clinical isolate of SCV2 USA-WA1/2020. Consistent with our results from C57BL/6 mice, only K18-hACE2 Tg mice whose lungs were pre-exposed to TLR ligands displayed a significant reduction in lung viral titers when assessed by either TCID_50_ or qPCR assays three days after infection **(Fig 2C).** This model is acutely susceptible to SCV2-induced disease, and we saw that reduced viral replication was reflected in significantly reduced SCV2-induced moribundity with CpG, but not Pm3 pre-treatment **(Fig 2D)**, despite no detectable differences in SCV2-induced lung pathology as determined by histological analysis three days after infection **(Fig S1C)**.

We next hypothesized that CpG- and Pm3-triggered protection after SCV2 infection may be associated with transcriptional changes in the lung. Three days after SCV2 infection we performed bulk RNA sequencing of K18-hACE2 Tg lungs that had been previously treated with CpG or Pm3. Gene Ontology (GO) enrichment analysis on differentially expressed genes (DEG) revealed genes involved in “antimicrobial peptide” responses for CpG pre-treated mice, while genes associated with “cellular interactions between lymphoid and non-lymphoid cells” were enriched in Pm3 pre-treated mice compared to SCV2 infected mice without prior TLR ligand exposure **(Fig 2E)**. The pathways shared by both CpG and Pm3 pre-treatments were associated with the activation of scavenger receptors, DAP12 signaling, and omega-3 and omega-6-derived bioactive lipid pathways **(Fig 2E).** Four significantly upregulated (*Fcrls, Ms4a7, Siglece, A930001A20Rik)* and three significantly downregulated (*Alox15, Marco, Mt2*) DEGs were shared between the SCV2 infected mice that were pre-treated with TLR agonists **(Fig 2F)**, which together reflect a transcriptional signature predominantly expressed in macrophages (ImmGen MyGeneSet) **(Fig 2G)**. It was notable that *Alox15*, which encodes the 12/15-lipoxygenase (12/15-LO) was found to be a significantly downregulated DEG. Lipoxygenases are key enzymes for the processing of omega-3 and omega-6-derived fatty acids into a variety of bioactive lipid mediators of inflammation and resolution ^54–56^. We hypothesized that TLR-induced downregulation of 12/15-LO in the lungs at the time of SCV2 exposure could decrease viral titers and, if so, 12/15-LO-deficient mice would accordingly display lower viral titers. However, when we infected *Alox15^−/−^* mice with SCV2 B.1.351, lung viral titers were instead significantly elevated in the knockout animals **(Fig 2H)**, suggesting that 12/15-LO deficiency does not license enhanced viral replication and instead is necessary for optimal SCV2 control. Together, these data revealed that the lungs of mice previously exposed to TLR-driven inflammation display relatively few transcriptional changes after SCV2 infection when compared to SCV2-infected lungs of mice not pre-treated with TLR agonists.

### Recent pulmonary TLR stimulation results in sustained tissue resident macrophage activation and inflammatory cytokines levels at the time of SCV2 exposure

To understand how the recent inflammatory history of the lung may condition the pulmonary microenvironment for improved viral control prior to SCV2 exposure, we analyzed lungs one week after TLR stimulation but before SCV2 infection. We started by asking whether improved viral control following recent local inflammation may simply be due to CpG or Pm3-mediated downregulation of the SCV2 entry receptor ACE2 prior to infection. We quantified ACE2 protein levels in lung homogenates of WT or K18-hACE2 Tg mice 7 – 10 days post-i.ph. treatment with TLR ligands and did not observe a reduction in the overall expression of either murine or human ACE2 protein **(Fig S2A)**. These data suggest that improved viral control in the lungs was not associated with a global reduction of ACE2 entry receptors at the time of SCV2 infection.

To further explore the heightened antiviral state after airway administration of TLR agonists, we performed bulk RNA sequencing 10 days following TLR activation of K18-hACE2 Tg lungs without SCV2 infection. In both CpG and Pm3 pre-treatment conditions, all significant DEGs were upregulated compared to RNA from control mice that received PBS **(Fig 3A)**. GO enrichment analysis revealed that similar pathways were enriched in CpG and Pm3-exposed lungs **(Fig 3B)**. GO terms that were shared between both protective conditions included those related to regulation of the complement cascade, chemokine receptors and chemokine signaling **(Fig 3B)**. Eleven significant DEGs (*2010008C14Rik, Aif1, C1qb, C3ar1, Calhm6, Ccr5, Cxcl9, Irgb10, Ms4a7, Saa3 and Trem2*) were identified in common between the two TLR-agonist treated groups compared to PBS controls **(Fig 3C).** Similar to the shared DEGs identified from lungs of mice that were TLR stimulated prior to SCV2 infection **(Fig 2G)**, shared DEGs after CpG or Pm3 lung administration were primarily expressed in macrophage subsets (ImmGen MyGeneSet) **(Fig 3C)**. Based on a chemokine and macrophage-enriched gene signature, we hypothesized that TLR-induced upregulation of CCR5, a chemokine receptor also expressed by macrophages and whose suppression has been identified as a genetic risk factor for COVID-19 ^12,21,57^, could account for the observed heightened antiviral state. When *Ccr5^−/−^*mice were infected with SCV2 B.1.35, lung viral titers were significantly elevated in the knockout animals compared to WT controls **(Fig 3D)**, supporting a critical role for CCR5 expression in SCV2 restriction, without prior TLR activation. However, CCR5 was dispensable for TLR agonist pre-treatment-mediated viral control, as *Ccr5^−/−^* mice still had reduced SCV2 viral loads after prior CpG or Pm3 exposure compared to untreated *Ccr5^−/−^* mice **(Fig 3E)**. Of note, although *Trem2* was a significantly upregulated DEG following TLR stimulation, Triggering Receptor Expressed on Myeloid cells 2 (TREM2) deficiency did not reverse the protective effect of CpG pre-treatment prior to SCV2 infection **(Fig S2B**) nor did deficiency in CCR2, another chemokine receptor highly expressed on monocytes and important for monocyte-derived macrophages **(Fig S2C)**.

**Figure 3:**
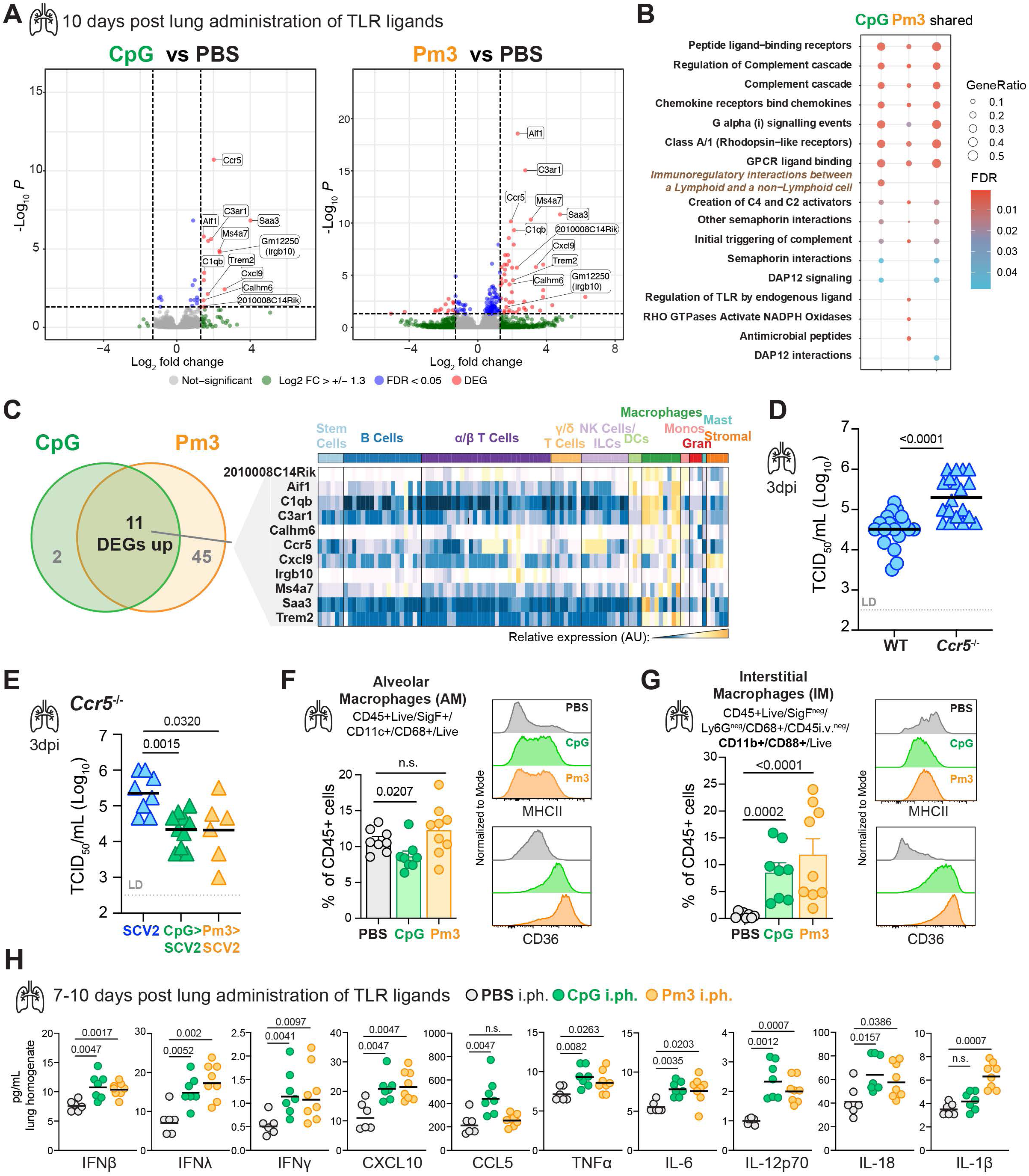
Recent pulmonary exposure to TLR agonists results in remodeling of the tissue-resident macrophage compartment and sustained inflammatory cytokine responses prior to SCV2 exposure. **(A – C)** K18-hACE2 Tg mice were i.ph. treated with PBS, 10µg CpG or 50µg Pm3. Ten days later, mice were euthanized, RNA was extracted from lung tissue and total RNA sequencing was performed; data is from 3-4 mice per group from one independent experiment **(A)** Candidate DEGs visualized by volcano plots comparing CpG- (left panel) or Pm3- (right panel) treated mice to the PBS control animals. DEGs upregulated and common to both treatment groups are labeled. **(B)** GO analysis of identified significant DEGs in the indicated groups compared to the PBS only controls. **(C)** Venn diagram showing the DEGs in common between the CpG- and Pm3-treated groups compared to PBS controls. The candidate DEGs were entered into ImmGen’s MyGeneset browser. Expression across cell types as analyzed by ImmGen are visualized in a heatmap, AU (arbitrary units), navy= lowest expression, orange= highest expression; ILCs (innate lymphoid cells), DCs (dendritic cells), Monos (monocytes), Grans (granulocytes), Mast (mast cells) **(D & E)** For the SCV2 (SCV2) infection, all mice were infected i.n. with SCV2 (B.1.351) and euthanized three days later as measured by TCID_50_, geometric mean, LD= limit of detection. **(D)** TCID_50_ viral loads in lungs of WT and *Ccr5*^−/−^ mice, n=19, data combined from five independent experiments. (E) *Ccr5*^−/−^ mice were given PBS, 10μg CpG or 50μg Pm3 one week before SCV2 infection with n=6-8, data combined from 2 independent experiments, geometric mean, LD= limit of detection. **(F)** Quantification of alveolar macrophages (AM) by flow cytometry as a percentage of CD45^+^ cells in whole lung from WT mice treated with PBS, 10μg CpG or 50μg Pm3 i.ph. one week prior and histograms depicting relative expression of MHC-II and CD36 **(G)** Quantification of CD11b^+^, CD88^+^ interstitial macrophages (IM) that are recruited into the lung parenchyma (intravascular CD45 negative (i.v^neg^)) by flow cytometry as a percentage of CD45^+^ cells in whole lung from WT mice treated with PBS, CpG or Pm3 i.ph. one week prior and histograms depicting relative expression of MHC-II and CD36, n= 8-9, data combined from two representative experiments, geometric mean and standard deviation. **(H)** Lungs were collected at seven and ten days after PBS, 10µg CpG or 50µg Pm3 i.ph. administration and homogenates were assayed for the indicated cytokines by multiplex bead array, n = 6-8, data combined from two experiments, geometric mean. **(D – H)** Statistical significances compared to PBS pretreated controls were determined by two-tailed Mann Whitney test.

Based on the transcriptional changes in innate and macrophage-associated genes, we hypothesized that prior pulmonary inflammation may remodel the lung microenvironment through the tissue-resident macrophage (TRM) compartment. In fact, TRMs have been shown to play important roles in SCV2 pathogenesis ranging from modulating lung-epithelial macrophage crosstalk, interferon responses, and antiviral T cell responses to contributing to pathology and cytokine storm at later disease stages ^58–63^. We, therefore, directly examined alveolar macrophages (AMs) **(Fig S3A)** and lung parenchymal residing interstitial macrophages (IMs) **(Fig S3B)** at the single cell level using multiparameter flow cytometry with intravascular (i.v.) staining to identify tissue-resident cells ^64^ one week after pulmonary CpG and Pm3 stimulation. In agreement with the macrophage-expressed genes identified in our transcriptional analysis we observed both quantitative and qualitative differences in AM and IM subsets associated with changes in lipid metabolism and alternative activation. While we saw a small but significant reduction of AMs after CpG, but not Pm3 exposure, the expression of class II major histocompatibility complex (MHCII) and CD36, a fatty acid translocase scavenger receptor ^65^, were increased after stimulation with both TLR agonists **(Fig 3F)**. In addition, expression of TREM2, TLR2, CD38 (ecto-NADase, activation and maturation marker ^66,67^, CD13 (aminopeptidase N, lung IM activation marker ^68^, ABCA1 (cholesterol efflux transporter), lectin-type oxidized LDL receptor 1 (LOX-1, scavenger receptor) and CD11b, all remained increased one week after either CpG or Pm3 stimulation in AMs **(Fig S3A)**. One week after treatment with TLR agonists we detected a significant population of CD88, CD11b+ IMs, characterized by low MHCII expression and high CD36 expression in CpG and Pm3-treated lungs that was largely absent in PBS control lungs **(Fig 3G**, **S3B)**. In addition to TREM2 and CCR5, this TRM population also expressed high levels of CD206, CD169, CD64, TLR2, CD38, LOX-1, ABCA1, CD14, and CD13 **(Fig S3B)**, indicative of increased fatty-acid oxidation and lipid catabolism in mature TRMs associated with tissue repair and homeostasis functions ^66,69,70^. In line with their phenotypic and metabolic changes, both AMs and IMs expressed increased levels of arginase-1 (Arg1) that persisted 7-10 days after one-time pulmonary TLR9 or TLR2 stimulation **(Fig S4A)**. Arginase and arginine metabolism have been implicated in COVID-19 severity, with studies supporting an immunosuppressive role for arginase-expressing myeloid cells in patients ^71–74^. However, arginase inhibition with Nω-hydroxy-nor-arginine (nor-NOHA) during SCV2 infection did not affect viral replication, nor did it reverse the protective lung conditioning by recent CpG or Pm3 exposure before SCV2 infection **(Fig S4B)**. Taken together, our data demonstrated that the pulmonary TRM compartment shows signs of remodeling and sustained activation alongside metabolic changes that are also reflected in total lung transcriptional profiling at the time of SCV2 infection.

Next, we screened lung homogenates from CpG or Pm3-treated mice by bead-based multiplex array for a variety of cytokines to better understand the inflammatory microenvironment of the lungs prior to SCV2 infection. Importantly, we detected a broad and persistent elevation of pulmonary cytokines even at 7 – 10 days after exposure to a single TLR ligand with marked increases in the interferons IFNβ, IFNγ, IFNλ alongside CXCL10, CCL5, TNFα, IL-6 and IL-12p70, IL-18 and IL-1 compared to PBS control mice **(Fig 3H)**. Taken together, these data indicate that recent inflammatory stimulation causes persistent perturbations in the lung microenvironment prior to SCV2 exposure that are associated with sustained alterations and activation of the TRM compartment as well as inflammatory cytokines, including antiviral IFNs.

### Pulmonary SCV2 replication is restricted by nucleic acid sensing and signaling by type I IFN and TNF, but not IL-1

Our transcriptional, cellular and cytokine profiling following recent airway TLR exposure of mouse lungs highlighted an expansive and persistent innate immune signature associated with antiviral conditioning of the lung microenvironment at the time of SCV2 infection. To better contextualize which innate immune pathways may be contributing to a heightened antiviral state before SCV2 infection, we next sought to determine the relative importance of various PRRs and key innate inflammatory cytokine pathways required for early control of SCV2 replication in the lung. Initial publications in mouse models reported conflicting results regarding the role of IFN-I in control of SCV2 viral replication ^3,56,75–82^. Therefore, we revisited IFN-I signaling in C57BL/6 mice infected with SCV2 B.1.351. We found that two different IFNAR1 deficient mouse strains had significantly increased viral titers as measured by TCID_50_ assay three days after exposure **(Fig 4A)**. Consistent with this, administering an anti-IFNAR1 neutralizing antibody one day prior to SCV2 infection also limited SCV2 replication **(Fig 4B)**. Measurements of SCV2 E sub-gRNA confirmed increased viral loads in all three IFNAR1-deficient models at this early timepoint after infection **(Fig S5A-B)**. Thus, consistent with the clinical evidence, initial SCV2 replication in the lungs of C57BL/6 mice is controlled via IFN-I-dependent pathways.

**Figure 4:**
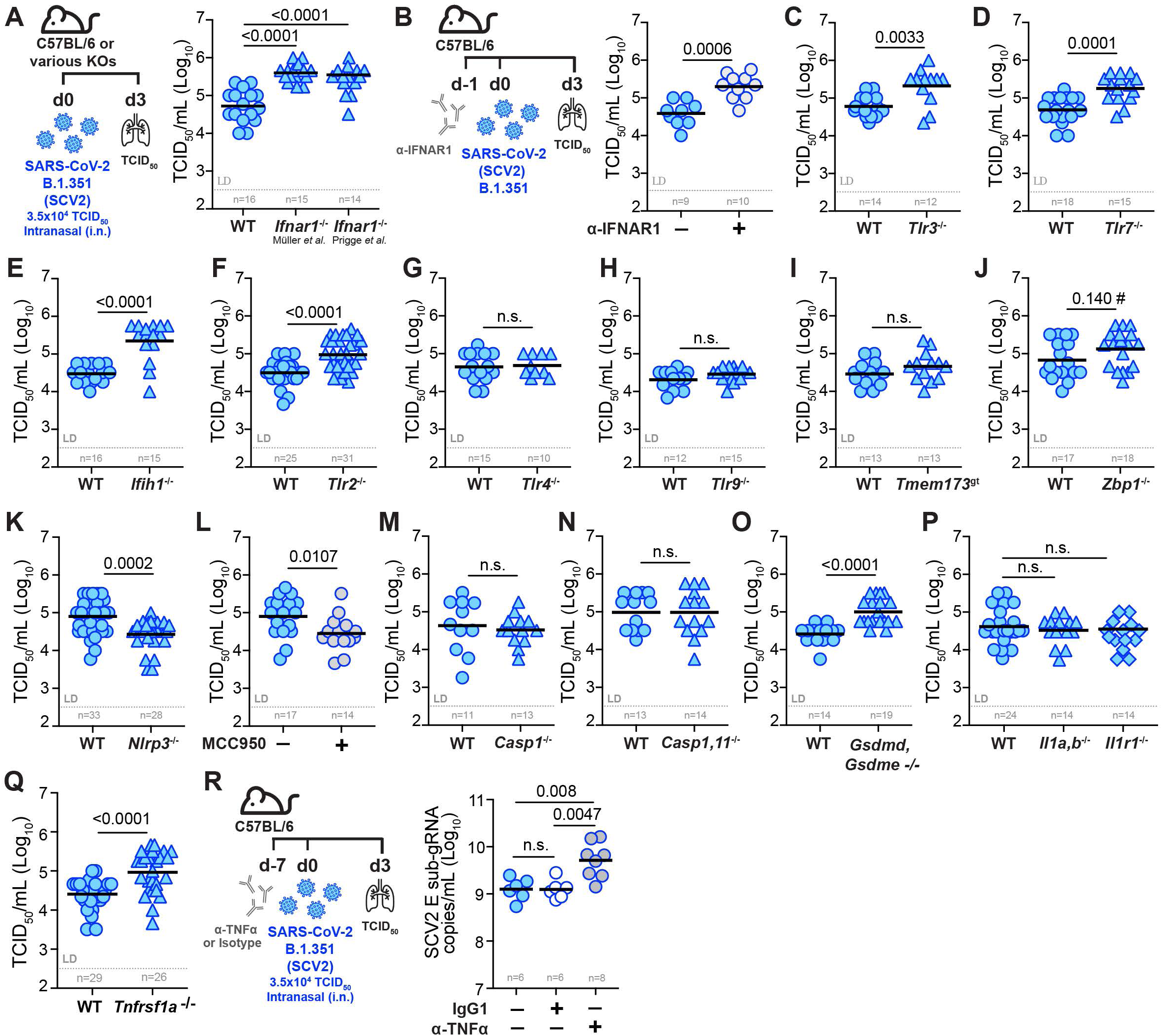
Pulmonary SCV2 replication is constrained by nucleic acid sensing, and signaling by IFNAR1 and TNFR1 but not IL-1R1. All mice were infected i.n. with 3.5×10^4^ TCID_50_ SCV2 (B.1.351) and euthanized three days later (d3). Viral loads were measured by TCID_50_ on Vero E6 cells except in (R). **(A)** Viral loads in lungs of WT and two different strains of *Ifnar1*^−/−^ mice. **(B)** Experimental set-up where WT mice were i.p. injected with a neutralizing anti-IFNAR1 monoclonal antibody one day before SCV2 infection and lung viral loads **(C – K)** Viral loads in lungs of WT and various PRR KO mice: **(C)** *Tlr3*^−/−^, **(D)** *Tlr7*^−/−^, **(E)** MDA5, *Ifih1*^−/−^, **(F)** *Tlr2*^−/−^, **(G)** *Tlr4*^−/−^, **(H)** *Tlr9*^−/−^, **(I)** *Tmem173*^gt^ (expresses an inactive variant of STING), **(J)** *Zbp1*^−/−^, **(K)** *Nlrp3*^−/−^. **(L)** Viral loads in lungs of WT mice that were i.p. injected with either PBS (-) or the NLRP3 inhibitor MCC950 (+) one day before and one day after SCV2 infection **(M – Q)** TCID_50_ viral loads in lungs of WT mice and mice deficient in inflammatory caspases or their substrates: **(M)** *Casp1*^−/−^ **(N)** *Casp1,11*^−/−^ **(O)** *Gsdmd,Gsdme*^−/−^ **(P)** *Il1a,b*^−/−^ and *Il1r1*^−/−^. **(Q)** Viral loads in lungs of WT mice and mice deficient in TNFR1, *Tnfrsf1a*^−/−^ **(R)** Schematic of experimental set-up where WT mice were i.p. injected with a neutralizing anti-TNFα monoclonal antibody seven days before SCV2 infection and viral loads in lung as measured by qPCR for the SCV2 E gene with and without anti-TNFα treatment. n indicated below each group, data combined from 2 – 6 independent experiments, geometric mean, statistical significance calculated by Mann Whitney test, LD= limit of detection, n.s.= not significant, #= indicates result that was not significant by TCID_50_ but showed a significant difference by qPCR (see **Fig S5E**).

Next, we examined the role of specific pattern recognition receptors (PRRs) and their relative importance in the innate restriction of pulmonary SCV2 replication. Consistent with what has been observed in patients with inborn errors of TLR3 or TLR7 ^2–4,17^ we found that mice deficient in TLR3 **(Fig 4C)** or TLR7 **(Fig 4D)** had significantly elevated lung SCV2 titers by TCID_50_ assay, as did mice deficient in the RNA-sensing cytoplasmic PRR melanoma differentiation-associated protein 5 (MDA5, encoded by *Ifih1*) **(Fig 4E)**. We next examined mice deficient in TLR2, a PRR reported to recognize the E and S protein of SCV2 ^83,84^. Lungs of *Tlr2*^−/−^ mice infected with SCV2 B.1.351 had significantly higher viral titers compared to WT mice **(Fig 4F)**, suggesting an important role for TLR2-triggered innate immunity in limiting early viral replication in mice. In contrast, neither TLR4 **(Fig 4G)** nor TLR9 **(Fig 4H)** were required for control of SCV2 replication in the lungs of mice three days after infection. Consistent with findings in *cGAS*^−/−^ mice ^85^, mice deficient in stimulator of IFN genes (STING, encoded by *Tmem173*) showed no difference in lung viral titers compared to WT controls **(Fig 4I)**. Mice lacking Z-DNA binding protein 1 (ZBP1) **(Fig S5C,D),** a nucleic acid sensor detecting both Z-configured DNA and RNA shown to play an important role in modulating IFN-I mediated lung inflammation in SCV2 at later stages ^86,87^, displayed a small yet significant increase in viral loads three days after infection when E sub-gRNA was measured by qPCR **(Fig S5E),** but when measured by TCID_50_ assay differences in viral titers did not reach statistical significance **(Fig 4J)**. Our data suggest, at best, a small effect of Z-formed nucleic acid sensing on viral replication, consistent with observations previously made in a SCV2 Ad-hACE2 mouse model^87,88^.

Because recent airway administration of TLR agonists resulted in elevated lung levels of IL-1β and IL-18, we next investigated whether inflammasome-related pathways contribute to innate control of viral replication in SCV2 infection. Indeed, both inflammasome activation and pyroptotic cell death have been implicated in SCV2 pathogenesis ^89–92^. Similar to observations made in an AAV-hACE2 mouse model of SCV2 infection ^93^, *Nlrp3^−/−^* mice infected with SCV2 B.1.351 exhibited significantly lower viral lung titers **(Fig 4K).** In addition, WT mice treated with the NLRP3 inhibitor MCC950 also had lower lung SCV2 titers **(Fig 4L)**. Next, we infected *Casp1^−/−^*and *Casp1,11^−/−^* mice to ask whether the observed decrease in viral loads in *Nlrp3^−/−^* mice was due to NLRP3-driven caspase-1 or caspase-11 activation. However, both *Casp1^−/−^*mice **(Fig 4M)**, as well as *Casp1,11^−/−^* mice **(Fig 4N)**, displayed similar lung SCV2 titers compared to WT mice. These data suggest that the inflammatory caspases −1 and −11 are dispensable for early viral control in the lungs of mice and unlikely operate downstream of NLRP3 to promote SCV2 replication in the first three days after infection. Interestingly, mice doubly-deficient in gasdermin-D and gasdermin-E (*Gsdmd,Gsdme*^−/−^) exhibited significantly increased lung SCV2 titers as measured by TCID_50_ assay **(Fig 4O)**, suggesting a key role for these terminal pore-forming proteins in the innate restriction of SCV2 viral titers in the mouse lung. As inflammasome activation and pore-formation are essential to both lytic pyroptotic cell death and the generation and secretion of leaderless cytokines like IL-1, we asked whether IL-1 signaling is required to control pulmonary SCV2 replication. Both *Il1a,b^−/−^*, and *Il1r1^−/−^* mice **(Fig 4P)** were able to control lung viral replication similarly to WT mice, arguing that IL-1R1-driven signals are dispensable for initial viral control in mice three days after infection with SCV2 B.1.351.

Finally, we investigated whether the pro-inflammatory cytokine TNFα was playing a role in modulating viral replication in the lungs early after SCV2 infection. TNFα is broadly inducible by diverse inflammatory cues and was also elevated in the lungs one week after TLR preconditioning. Of note, TNFR1 deficient (*Tnfrsf1a*^−/−^) mice infected with SCV2 B.1.351, displayed defective control of viral replication as quantified by TCID_50_ assay **(Fig 4Q)** or by qPCR for E sub-gRNA **(Fig S5F).** TNFα neutralization by monoclonal antibody treatment prior to SCV2 infection likewise resulted in elevated lung viral burdens compared to IgG1 control-treated mice **(Fig 4R)**. Therefore, our data demonstrate that TNFR1-mediated pro-inflammatory signals contribute to limiting early SCV2 replication in the lung. Taken together, our survey of innate inflammatory pathways relevant for initial viral control not only revealed an expected important role for IFN-I but also delineated key PRRs and pro-inflammatory pathways important for limiting pulmonary viral replication prior to the onset of adaptive SCV2-specific responses that may contribute to the establishment of an antiviral lung microenvironment.

### Recent IFN-I-dependent and -independent inflammatory conditioning of the lung promotes suppression of viral replication

Because we detected elevated levels of interferons **(Fig 3H)** after TLR agonist conditioning and the transcriptional pathway analyses highlighted “immunoregulatory interactions between a Lymphoid and a non-Lymphoid cells” **(Fig 2E**, **3B)** we next hypothesized that the inflammatory changes in the lung microenvironment prior to SCV2 infection might also extend to changes in lung epithelial cells (ECs), the primary target and reservoir of SCV2 replication ^94,95^. Thus, we next examined mouse pulmonary EC subsets prior to SCV2 infection after conditioning with the different protective pulmonary inflammatory stimuli described above, ranging from sterile inflammation to bacterial and viral infection. Using flow cytometry we assessed CD45^neg^ CD31^neg^ CD326^+^ pulmonary EC subsets in the mouse lung, including CD24^neg^ Podoplanin (PDPN)^hi^ MHC-II^low^ type I alveolar epithelial cells (AECIs), CD24^neg^ PDPN^low^ MHC-II^hi^ type II alveolar epithelial cells (AECIIs), CD24^neg^ PDPN^neg^ MHC-II^neg^ ECs (Other ECs), CD24^+^ CD49f^hi^ Bronchiolar Epithelial Cells (BECs) and CD24^+^ CD49f^low^ club cells **(Fig S6A)**. We measured the IFN-activation of ECs by quantifying the expression of the IFN-inducible surface markers (ISMs) CD317 (BST2, HM1.24, PDCA1, tetherin) and Sca-1(Ly6A/E) by flow cytometry ^40,96,97^ **(Fig 5A**, **S6B)**. All inflammatory and infectious stimuli induced changes in ISM upregulation across all cell types, with Pm3 providing the weakest EC ISM activation and, expectedly, chronic *Mtb* infection the strongest **(Fig 5A**, **S6B).** One week after pulmonary TLR agonist administration, only Sca-1 on BECs was significantly elevated with Pm3, while CpG increased both CD317 and Sca-1 across most EC subsets, supporting more potent IFN-induced conditioning of the pulmonary epithelium following activation of TLR9 compared to TLR1/2 **(Fig 5A**, **S6B)**. Additionally, 30 days following IAV infection, we detected increased expression of Sca-1, but not CD317, on all ECs except the Other ECs group, which had significantly decreased expression of both ISMs one month after IAV infection **(Fig 5A**, **S6B)**, agreeing with previously published data showing that IAV does not induce CD317 on lung ECs ^98^. Three days after pulmonary *S. aureus* infection, both ISMs were upregulated on BEC and club cells with the highest CD317 activation in AECIIs and BECs compared to naïve control animals **(Fig 5A**, **S6B)**. Notably, five days after pulmonary OVA administration, CD317 was very highly elevated on AECIIs and BECs in the OVA/Alum model of allergic asthma **(Fig 5A**, **S6B)**, suggesting IFN activation of ECs even in a type-II-response-dominated inflammatory setting. Taken together, all tested inflammatory and infectious settings led to heightened IFN-activation of ECs prior to SCV2 exposure when compared to naïve animals, suggesting the possibility that inflammatory conditioning of the lung microenvironment involves direct sensitization of ECs for enhanced control of subsequent SCV2 infection.

**Figure 5:**
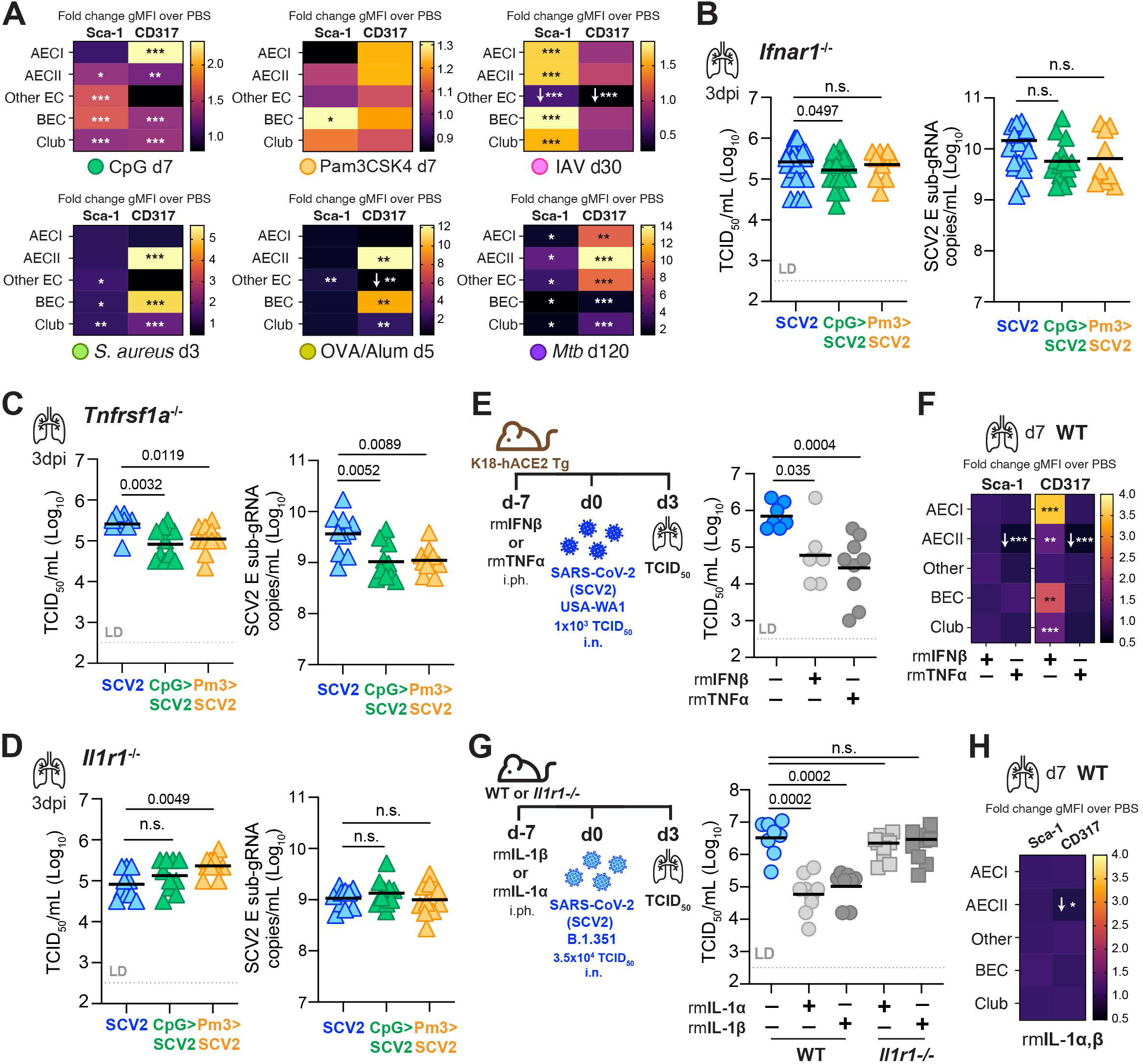
Recent IFN-I dependent and -independent inflammatory conditioning of the lung promotes SCV2 replication control at the tissue level. **(A)** Heatmap of fold change in the geometric mean fluorescence intensity (gMFI) of IFN-inducible surface marker (ISM) expression of Sca-1 and CD317 measured by flow cytometry on lung epithelial cell (EC) subsets from lungs of mice treated with various inflammatory or infectious stimuli compared to those from PBS control animals at the indicated time points without SCV2 infection, n=5-14, data is pooled from 2 – 4 independent experiments, for all conditions except OVA/Alum which was done once. **(B)** *Ifnar1*^−/−^, **(C)** *Tnfrsf1a*^−/−^, or **(D)** *Il1r1*^−/−^ mice were i.ph. treated with PBS, 10µg CpG or 50µg Pm3 one week before SCV2 (B.1.351) infection. Viral loads in lungs were quantified by TCID_50_ or sub-gRNA SCV2 E qPCR, n=10-26, data are combined from 2 – 3 independent experiments each **(E)** Schematic of K18-hACE2 Tg mice administered 5μg recombinant mouse TNFα (rmTNFα) or 2.0×10^4^U recombinant mouse IFNβ (rmIFNβ) once, one week before infection with SCV2 (USA-WA1/2020) and lung viral titers, n=7-10, two independent experiments **(F)** Fold change in gMFI of Sca-1 and CD317 measured by flow cytometry on lung EC subsets of mice treated one week prior with rmTNFα or rmIFNβ, data is pooled from three independent experiments, n= 15-22, *= p<0.05, **=p<0.01, ***=p<0.001, if not indicated, differences were not significant. **(G)** WT or *Il1r1*^−/−^ mice given 200U recombinant mouse IL-1α (rmIL-1α) or IL-1β (rmIL-1β) i.ph. one week before SCV2 (B.1.351) infection and ung viral titers, n=8-10, data are combined from two independent experiments. **(H)** Fold change in gMFI of Sca-1 and CD317 measured by flow cytometry on EC subsets from lungs of mice treated one week prior with 200U rmIL-1α + rmIL-1β (100U each), n= 9-10, data is pooled from two independent experiments. Geometric mean, statistical significance determined by two-tailed Mann-Whitney test, LD= limit of detection, *= p<0.05, **=p<0.01, ***=p<0.001, n.s.= not significant, if not indicated differences were not significant, white arrows indicate direction of fold change.

Since IFNs were elevated and we observed increased expression of IFN-inducible markers on lung ECs one week post pulmonary treatment with TLR-agonists, we next tested whether this elevated baseline IFN-I signaling at the time of SCV2 exposure was sufficient for the viral control induced by prior CpG or Pm3 exposure. Indeed, in contrast to WT animals **(Fig 2A**, **S1A)**, *Ifnar1^−/−^*animals failed to display significantly reduced SCV2 viral titers after recent pulmonary CpG or Pm3 treatment when measured by TCID_50_ or qPCR **(Fig 5B)**. These data support the hypothesis that recent IFN-I-driven changes in the lung microenvironment may be sufficient to condition the lung for enhanced viral restriction when exposed to SCV2. Remarkably, Pm3-mediated protection was abrogated in the absence of *Ifnar1*, despite being a more potent inducer of NF-κB than CpG and only very weakly sustaining Sca-1 expression in BECs **(Fig 5A)**. Because we observed increased levels of TNFα in the lungs after CpG or Pm3 treatment **(Fig 3H)** and a requirement for TNFR1 in initial viral control after SCV2 infection **(Fig 4Q, 4R)**, we next asked whether the sustained increase in TNFα seen after TLR conditioning was contributing to the heightened antiviral state, similar to what we observed with IFN-I. However, when we exposed TNFR1 deficient (*Tnfrsf1a*^−/−^) mice to CpG or Pm3 one week prior to infection with SCV2 B.1.351, the ability of CpG or Pm3 to significantly suppress pulmonary SCV2 loads was retained in *Tnfrsf1a*^−/−^ animals **(Fig 5C)**. Thus, while TNFα appears to be critical for viral control after SCV2 infection, it is dispensable for TLR9 or TLR1/2-mediated enhanced antiviral conditioning of the lungs. Importantly, although IL-1R1 was not required for control of SCV2 **(Fig 4P)**, IL-1β remained elevated in lung homogenates one week after TLR1/2 stimulation **(Fig 3H)**. We, therefore, tested the contribution of IL-1 to enhanced viral control following CpG or Pm3 exposure and found that unlike WT and like *Ifnar1^−/^ mice*, *Il1r1^−/−^* mice failed to restrict SCV2 replication when measured by either TCID_50_ or qPCR **(Fig 5D)**. These data suggest a requirement for effective signaling through either IFNAR1 or IL-1R1, but not TNFR1 to effectively suppress SCV2 following TLR9 or TLR1/2 conditioning. Moreover, while without recent exposure to inflammatory stimuli or other pathogens, IL-1R1 was dispensable for viral control after SCV2 infection, inflammatory conditioning via TLRs was nonetheless able to co-opt the IL-1 pathway to promote a heightened antiviral state in the lungs of mice.

Lastly, to further de-construct how diverse inflammatory stimuli may condition the lung microenvironment to improve early viral control, we asked whether exposure to inflammatory cytokines themselves is sufficient. Based on the ISM profiling results of lung EC subsets, we hypothesized that IFNs are likely variably expressed amongst the diverse settings of lung perturbations examined, while TNFα is likely more broadly induced across a wide range of inflammatory stimuli. Thus, to determine whether recent activation of IFNAR1 or TNFR1 signaling pathways is sufficient to create a heightened antiviral state and effectively restrict SVC2 replication in the mouse lung, recombinant IFNβ, or TNFα were given to K18-hACE2 Tg mice by i.ph. seven days prior to infection with SCV2 USA-WA1/2020. Notably, both IFNβ and TNFα were able to significantly reduce the SCV2 burden as measured by TCID_50_ assay at three days post infection by ~1.5 logs **(Fig 5E)**. Moreover, when we profiled ISM expression on ECs in WT mice one week after receiving pulmonary stimulation with recombinant IFNβ, or TNFα, only mice that received IFNβ, but not TNFα, showed significant upregulation of CD317 **(Fig 5F)**, despite equal suppression of viral replication. This finding demonstrates that CD317 ISM upregulation is specific to IFNs and that TNFα-mediated antiviral protection occurs independently of upregulation of ISM in ECs measured here. Thus, in addition to the known antiviral effects of IFNβ, our data reveal that increased TNFα driven inflammation at the time of SCV2 exposure is potently able to confer a heightened antiviral state in the lung. Finally, we asked whether IL-1α and IL-1β are equally able to provide antiviral inflammatory conditioning of the lung. Indeed, pulmonary administration of either IL-1α or IL-1β one week prior to SCV2 infection of C57BL/6 mice, was able to significantly enhance early viral control in the lungs of WT but not *Il1r1^−/−^* animals as measured by TCID_50_ **(Fig 5G)**. Similar to TNFα administration, IL-1 preconditioning of the lung did not result in indirect activation of IFN pathways in the pulmonary epithelium, as we failed to detect ISM marker expression on different lung EC subsets prior to infection with SCV2 **(Fig 5H)**. In summary, the above data revealed that both IFN-I-dependent and -independent pro-inflammatory pathways can promote an effective, antiviral inflammatory tone in the mouse lung and suggest that prior engagement of the IL-1 or TNFα signaling pathways is sufficient to restrict pulmonary SCV2 replication in mice. Taken together our findings provide a molecular framework and *in vivo* evidence that immunologically diverse pulmonary exposure histories, including those that only modestly trigger IFN responses, can potently impact initial pulmonary SCV2 replication.

## DISCUSSION

Here, we establish that recent or ongoing pulmonary inflammatory stimuli, such as newly resolved respiratory infections, sterile allergic inflammation, or TLR agonist and cytokine-induced responses, modulate the early antiviral response in the lungs upon SCV2 encounter. We demonstrate that elevated baseline induction of pro-inflammatory TNFα or IL-1 responses, in addition to IFN-I, impart antiviral activities capable of lowering initial viral titers. Moreover, our findings suggest that prior engagement of the IL-1 or TNFα signaling pathways can restrict SCV2 replication in the mouse lung.

IFNs play critical roles in limiting viral replication early after SCV2 infection, yet at later stages of infection and when dysregulated, they also contribute to disease ^8,81,85,99–103^. Our study is centered around innate factors limiting viral replication in the lung prior to the onset of adaptive immunity. Most of the inflammatory pathways we show here as required for early innate control of viral titers have been implicated in disease progression and mortality at later stages and have, therefore, contextual roles depending on disease stage. For example, while the SCV2 spike and envelope proteins trigger inflammation through the binding of TLR2 and blockade of TLR2 signaling extended survival in mice ^83,84^, we demonstrate that TLR2 is required for optimal control of viral replication in the lungs of mice early after infection. Similarly, TNFα, IL-1 and inflammasome pathways have been implicated in the deleterious effects of cytokine storms and cell death later in disease ^89,92,104–106^. We found that the pro-inflammatory TNFR1 pathway is critical for early viral control of SCV2 replication in mice and that pulmonary preconditioning with recombinant TNFα potently limits SCV2 viral titers in the lung. Of note, patients undergoing TNFα blockade as a long-term treatment for dermatological or rheumatic diseases showed no increase in hospitalizations or severe COVID-19 disease ^107–109^. However, the impact on early control of viral replication is difficult to ascertain from such clinical studies. In regard to IL-1 and inflammasomes, our data agree with prior findings of decreased viral titers early after infection in *Nlrp3^−/−^*mice ^93^, supporting a negative role for NLRP3 activation at both early and later stages of SCV2 infection. However, we also provide evidence that uncouples NLRP3 from its known downstream effector proteins as we show here that early viral titers were unchanged in *Casp1^−/−^*, *Casp1,11*^−/−^, *Il1r1^−/−^* and *Il1a,b^−/−^* animals and were in fact higher in *Gsdmd,Gsdme^−/−^* mice. Further studies will be required to address the differential roles revealed here for gasdermins and NLRP3 in early innate mucosal immunity to SCV2 compared to later stages of disease. Moreover, although the IL-1R1 signaling pathway was not required to control early viral replication during SCV2 infection, pulmonary IL-1-driven inflammatory preconditioning prior to SCV2 infection established an antiviral state, like TNFα and IFNβ preconditioning. Thus, TNFα and IL-1, in addition to IFNs, are important for viral control in the lung around the time of SCV2 exposure, while at later stages their release must be controlled to prevent systemic cytokine storm and pathogenic inflammation.

While an important conclusion from the present study is that diverse infectious and sterile inflammatory stimuli can precondition the lung for enhanced early innate control of SCV2 replication, the precise molecular and cellular mechanisms underlying enhanced viral control are most likely as varied as the stimuli used. Future studies will be required to carefully delineate cellular and molecular mechanisms *in vivo* for each inflammatory scenario. By narrowing from complex pathogens to TLR ligands and individual cytokine responses we can, however, speculate on some general features that may be common among those stimuli and induce an innately protective antiviral state in the lungs of mice. First, it is conceivable that distinct inflammatory axes converge on the induction of antiviral IFNs and that broad IFN-driven responses are sufficient and necessary for enhanced antiviral immunity in the lung. Indeed, pulmonary administration of IFN-I (IFNα ^102^ or IFNβ as shown here), IFN-II (IFNγ ^40^) or IFN-III (IFNλ ^85,110,111^) prior to infection all lowered SCV2 viral titers in the lungs of mice. Pulmonary delivery of the STING agonist 2’3’ cGAMP ^112^, the TLR3/MDA5 agonist Poly (I:C) ^113^ or the RIG-I agonist SLR14 ^114^ were also shown to effectively reduce lung viral SCV2 burden when given prophylactically. In addition, IFN-I and IFN-III dependent control of viral replication in lung ECs has been shown to affect disease trajectories prior to the onset of T cell-mediated control ^8,15,103,115^. In fact, SCV2 replication levels are associated with the likelihood of viral transmission, and the ability of children to mount a more robust protective innate immune response compared to adults correlates with reduced viral replication in ECs ^5,6,116–119^. We provide evidence that the diverse inflammatory stimuli used in the current study, apart from recombinant TNFα and IL-1 administrations, all promoted IFN-driven activation of lung ECs as we observed increased expression of the ISMs Sca-1 or CD317 on various ECs in most settings prior to SCV2 infection. The pro-inflammatory cytokines IL-1 and TNFα can stimulate IRF-1 and IRF-3, transcriptional activators of chemokines and IFN-Is, thus promoting antiviral responses indirectly via IFN induction ^120–124^. Nevertheless, any potential indirect up-regulation of IFN-I by recombinant TNFα or IL-1 was insufficient to cause lung EC ISM expression when compared to recombinant IFNβ administration itself. While TNFα and IL-1 administration failed to upregulate Sca-1 or CD317 on ECs, it is possible that ISMs not examined in this study could have been activated indirectly by TNFα or IL-1. Additionally, the infectious and inflammatory stimuli used in our study are known to result in changes to the TRM compartment ^70^, and we show here that recent one-time TLR preconditioning potently remodeled the TRM compartment towards a metabolically altered tissue repair phenotype with increased arginase-1 expression. However, when we functionally inhibited arginase or tested IM involvement via CCR5, CCR2, or Trem2 deficient mice, TLR agonist-mediated protection was not abrogated, suggesting that arginase or IMs may not be sufficient to create the observed heightened antiviral state. We have also not ruled out that AMs may produce IFN-I in response to TLR stimulation, as reported during infections with other respiratory RNA viruses ^125,126^, or that lung ECs directly respond to CpG or Pm3 to generate IFNs and limit viral replication. Thus, direct or indirect activation of broad IFN responses close to the time of SCV2 exposure may be sufficient to enhance early innate viral resistance.

A second possibility is that TNFα and IL-1 exert IFN-independent antiviral effector functions ^127–130^. TNFα induces pro-inflammatory responses through TNFR1 complex I and noncanonical NF-κΒ activation as well as modulating cell death via complex II ^131^. Although excessive inflammatory cell death has been implicated in severe COVID-19 disease ^132,133^, killing infected cells and modulating cell death also represents a central antiviral strategy. TNFα can induce both apoptosis through caspase-8 and necroptosis, which utilizes the pseudokinase Mixed Lineage Kinase Domain-Like (MLKL). While we report here increased viral titers in TNFR1 deficient mice, mice lacking caspase-8 and/or MLKL had similar SCV2 viral titers when compared to lungs of WT animals ^134,135^. Besides modulation of cell death, TNFα drives changes in intracellular metabolism, including glycolysis, shown to be important for its cell-intrinsic antiviral activities ^127^. Both TNFα and IL-1 are also strong inducers of inflammatory chemokines and early and effective recruitment of innate immune cells has emerged as an important factor for viral replication in mice ^136^, while risk for severe COVID-19 has been linked to genetic regions expressing multiple chemokines ^12,21,57^. Thus, future studies will need to thoroughly delineate IFN-dependent and independent antiviral effects of TNFα and IL-1 during both antiviral preconditioning of the lung as well as in response to SCV2 infection. These effects may include initial TRM-EC interactions, the contribution of epigenetic modifications in TRMs and ECs, early cell death events, activation of AMs, induction of chemokines required for IM recruitment, or cell-intrinsic effects within lung EC subsets.

Nonhuman primates infected with SCV2, despite being genetically diverse and outbred, similar to the human population, present with only very mild disease associated with control of viral replication prior to antigen-specific T-cell responses ^137,138^. Importantly, nonhuman primates are typically not housed under abnormally hygienic specific pathogen-free (SPF) conditions, like most experimental mice, and have experienced diverse infectious immunological stimuli. In contrast to SPF mice, feral and pet-store mice that have been microbially exposed to naturally occurring infections exhibit elevated IFN and inflammatory responses and mount more human-like responses, resulting in increased viral control compared to SPF mice ^139–141^. Moreover, sequential infection of SPF mice can recapitulate some aspects of the naturally occurring prior exposure histories in wild mice and similarly promotes more human-like inflammatory immune responses ^142^. Our findings here are consistent with these prior observations on how immunological exposure history can shape the outcome of subsequent infections and we propose that feral or pet-store mice, like the preconditioned mice used here, would display enhanced SCV2 viral control compared to SPF mice, as has already been reported for infection with Lymphocytic Choriomeningitis Virus ^139^.

Our findings may also help provide an immunological basis for certain clinical observations in specific patient populations. For example, children are among the most widely infection-exposed patient populations, and most children have milder SCV2 infection outcomes compared to adults or the elderly. Besides age itself, one additional factor contributing to milder outcomes may also be recent infectious exposure histories. The concept of ‘immune debt’ and ‘immunity gap’ ^143–146^ suggests that during the pandemic, the unprecedented non-pharmaceutical interventions, including lock-down and masking, led to a significant decrease in exposure to common respiratory childhood diseases. After restrictions were lifted, however, a dramatic surge in pediatric respiratory disease was observed, arguing that the lack of pulmonary immune stimulation made children more vulnerable to community-acquired infections ^146–150^. Our study provides experimental evidence that diverse recent pulmonary exposures can indeed significantly impact subsequent innate viral infection control in the lung.

Children and adults with asthmatic diseases were initially thought to have more severe COVID-19 outcomes, given known deficits in antiviral immunity and that common respiratory viruses can exacerbate asthma. However, many clinical studies failed to show an expected increase in the prevalence of asthmatic patients among COVID-19-infected individuals and instead concluded that the relative risk of severe COVID-19 was relatively small ^32,34,151–153^. We show here in a murine experimental OVA/Alum asthma model that underlying allergic-type II-driven inflammation at the time of SCV2 exposure significantly enhances innate viral replication control in the lungs, arguing that innate aspects of type II immune responses might provide potential antiviral protective rather than detrimental effects early during SCV2 infection. Delineating the precise role type II associated cytokine, chemokines, and cell types promoting innate early control of SARS-CoV-2 replication will provide important mechanistic insight and context for clinical studies where type-2 associated immune responses were shown to either positively or negatively associate with COVID-19 outcomes ^151,154^.

There was also increased concern for patients with CF, an inherited, autosomal recessive disorder caused by mutations in the gene encoding the anion channel Cystic Fibrosis Transmembrane Conductance Regulator (CFTR) that can lead to chronic pulmonary infections and respiratory failure. However, COVID-19 incidence estimates in CF were reported to be lower than in the general population with often less severe outcomes than originally anticipated ^36,155,156^. In our experimental mouse model, we demonstrate that recent pulmonary infection with *S. aureus* resulted in significantly enhanced innate control of SCV2 replication. The most common lung pathogens that colonize CF patient include *Pseudomonas aeruginosa* and *S. aureus*, including methicillin-resistant *S. aureus* (MRSA), and *Aspergillus* ^157^. Therefore, it may be worth exploring whether bacterial colonization status at the time of SARS-CoV2 exposure is a contributing factor to the clinical outcome of COVID-19 in CF patients.

In conclusion, our study provides a foundational experimental framework together with *in vivo* evidence that immunologically diverse pulmonary exposure histories, including those that only modestly trigger IFN responses, can potently impact initial pulmonary SCV2 replication. Our findings open up the intriguing possibility that the recent exposure history and the inflammatory microenvironment of the lung proximal to the time of SCV2 exposure may be a significant factor contributing to the diverse clinical outcomes seen in people with COVID-19.

## MATERIALS & METHODS

### Mice

*Tlr4*^−/−^ mice (B6(Cg)-*Tlr4^tm1.2Karp^*/J; JAX #29015), *Tlr7*^−/−^ mice (B6.129S1-*Tlr7^tm1Flv^*/J; JAX #3080) ^158^, K18-hACE2 hemizygous transgenic mice (B6.Cg-Tg(K18-ACE2)2^Prlmn/J^; JAX #34860) ^159^, *Tmem173*^gt^ I199N mutant mice (C57BL/6J-*Sting1^gt^*/J; JAX #17537) ^160^, *Trem2*^−/−^ mice (C57BL/6J-*Trem2^em2Adiuj^*/J; JAX #27197) ^161^, *Casp1*^−/−^ mice (B6.Cg-*Casp1^em1Vnce^*/J; JAX #32662) ^162^, *Alox15*^−/−^ mice (B6.129S2-*Alox15^tm1Fun^*/J; JAX #2778) ^163^, and *Ifnar1*^−/−^ mice (B6(Cg)-Ifnar1^tm1.2Ees^/J; JAX stock #28288) ^164^, were purchased from Jackson Laboratories (Bar Harbor, ME). *Tlr2*^−/−^ mice ^165^, *Il1a,b*^−/−^ mice ^166^, *Tlr9*^−/−^ mice ^167^ were previously described. *Gsdmd,Gsdme*^−/−^ mice ^168^ were kind gifts of Dr. Feng Shao (NIBS, China). C57BL/6 mice (Taconic farms), C57BL/6 mice expressing a *Foxp3*-GFP reporter (C57BL/6-*Foxp3^tm1Kuch^*) ^169^ or the Thy1.1 allele (B6.PL-Thy1^a^/CyJ) were used as wild type C57BL/6 controls in experiments. *Foxp3*-GFP mice, Thy1.1 mice, *Ifnar1*^−/−^ mice (B6.129S2-*Ifnar1^tm1Agt^* backcrossed to B6 for 12 generations) ^170^, *Tlr3*^−/−^ mice (B6;129S1-*Tlr3^tm1Flv^*/J backcrossed to B6 for 11 generations) ^171^, *Ifih1*^−/−^ mice (B6.Cg-*Ifih1^tm1.1Cln^*/J) ^172^, *Ccr2*^−/−^ mice (B6.129S4-*Ccr2^tm1Ifc^*/J) ^173^, *Nlrp3*^−/−^ mice (B6N.129-*Nlrp3^tm2Hhf^*/J) ^174^, *Tnfrsf1a^−/−^* mice (C57BL/6-*Tnfrsf1a*^tm1Imx^/J) ^175^, *Ccr5*^−/−^ mice (B6.129P2-*Ccr5^tm1Kuz^*/J) ^176^, *Casp1,11*^−/−^ mice (B6N.129S2-*Casp1^tm1Flv^*/J) ^177^, *Il1r1*^−/−^ mice (B6;129S1-*Il1r1^tm1Rom^l*/J backcrossed to B6 for 12 generations) ^178^ were all obtained through a supply breeding contract between NIAID and Taconic Farms. *Zbp1^−/−^*mice were made in-house by CRISPR/Cas9 genetic targeting as detailed below. Both male and female mice, 8-16 weeks old, were used in experiments and all mice within individual experiments were age and sex matched. Genotyping was performed by Transnetyx using real-time PCR and genetic background analysis was submitted through Transnetyx and performed by Neogen using the MiniMUGA platform to confirm that all mice were on a C57BL/6 background. All animals were bred and maintained in an AAALAC-accredited ABSL2 or ABSL3 facility at the NIH and experiments were performed in compliance with an animal study proposal approved by the NIAID Animal Care and Use Committee.

### Generation of *Zbp1*^−/−^ mice

*Zbp1*^−/−^ mice were made by the NIAID Mouse Genetics and Gene Modification (MGGM) Section by microinjection of *Cas9* mRNA and the following guides: 5’ sgRNA GTTTCCGGGATGGTAACAGC and 3’ sgRNA CTGGGACCCACGCGAGGTGA into mouse embryos resulting in deletion of exon 1 to create null allele *Zbp1* c.-119_34+22del **(Fig S1C)**. G0 was crossed to C57Bl/6NTac mice to isolate the null allele in G1, which were intercrossed to homozygosity (screened using genotyping primers fwd: TCAGATAGAGCTCTCCCGGT, rev: TAGACAGGGTATGTAGTCTCAGC). Zbp1 knockout was validated at the protein level by western blotting of lysates from bone marrow-derived macrophages (BMDMs, differentiated for seven days with 50ng/mL M-CSF) and adherent peritoneal exudate cells (PECs) incubated with and without 200ng/mL lipopolysaccharide (LPS, InvivoGen, #tlrlpb5lps) for 6 hours **(Fig S1D)**. Cells were lysed in RIPA buffer and denatured by boiling in a final concentration of 70 mM SDS. Lysate from 5.6×10^4^ cells per lane was separated by SDS-PAGE and transferred to 0.2 µm nitrocellulose membranes. ZBP1 was detected using mouse-α-ZBP1 (Adipogen, #AG-20B-0010-C100, 1:1000) and donkey-α-mouse-HRP (Jackson ImmunoResearch, #715-035-150, 1:10000), as a positive control for LPS stimulation, pro-IL-1β was detected using goat-α-IL-1β (R&D, #AB-401-NA, 1:1500) and bovine-α-goat-HRP (Jackson ImmunoResearch, #805-035-180, 1:10,000) and actin was detected using mouse-α-actin-HRP (Santa Cruz, #sc-47778, 1:5000).

### SCV2 infections

SCV2 hCoV-19/USA-WA1/2020 (Pango lineage A, GISAID reference: EPI_ISL_404895.2) (USA-WA1/2020) and SCV2/human/ZAF/KRISP-K005325/2020 beta variant of concern (Pango lineage B.1.351, GISAID reference: EPI_ISL_678615) (B.1.351) were obtained from BEI resources (NIAID, NIH). Viral stocks were generated and sequenced as previously described ^39,179^. Mice were anesthetized with isoflurane and infected i.n. with 35μL inoculum containing 1.0×10^3^ TCID_50_ USA-WA1/2020 or 3.5×10^4^ TCID_50_ B.1.351. Inoculum was quantified by TCID_50_ assay in Vero E6 cells (American Type Culture Collection, #CRL-1586).

### *Mtb* infection of mice

Mice were infected with *Mtb* H37Rv-mCherry (50 – 200 CFU) by aerosol using a Glas-Col whole-body inhalation exposure system as previously described ^180^. Mice were infected with SCV2 75 – 100 days post *Mtb* infection.

### *Staphylococcus aureus* infection of mice

Mice were anesthetized with isoflurane and infected i.ph. with 5.6×10^7^ CFU *S. aureus* (USA300). Pulmonary delivery doses were confirmed by plating inocula on brain-heart infusion agar (BD Biosciences, #241830) and incubating at 37°C overnight. Mice were infected with SCV2 three days post *S. aureus* infection.

### Influenza A virus H1N1 infections

Mice were anesthetized with isoflurane and infected i.n. with 500 TCID_50_ Influenza A virus (IAV; A/Puerto Rico/8/34, H1N1 [PR8]) ^181^. Mice were then infected with SCV2 30 days post IAV infection.

### OVA-Alum lung allergy model

Mice were injected intraperitoneally (i.p.) twice, 14 days apart, with 100µg Ovalbumin (OVA, Sigma-Aldrich, #A5503) in 200µl containing 12.5% Imject^TM^ alum adjuvant (ThermoFisher, #77161). Ten days after the last injection mice were anaesthetized with isoflurane and challenged i.n. with 30µg OVA in 30µL injection grade sterile saline. Mice were infected with SCV2 5 – 6 days after i.n. OVA challenge.

### Treatment of mice with TLR agonists, recombinant cytokines, inhibitors or neutralizing antibodies

Mice were anesthetized with isoflurane and treated i.ph. with 30 – 50μL injection-grade saline containing TLR agonists (10μg CpG ODN 2088, CpG type B, Invivogen, #tlrl-1826; 50μg Pam3CSK4 (Pm3), Invivogen, #vac-pms) or recombinant cytokines (5µg TNFα, PeproTech, #315-01A; 2.0×10^4^U IFNβ, PBL, #12400-1; 200U IL-1α #211-11A, 200U IL-1β #211-11Β or both together at 200U total, PeproTech) to allow for pulmonary delivery one week (unless otherwise stated in the figure legends) prior to SCV2 infection. For neutralization of cytokine signaling, mice were i.p. injected with 500μg anti-TNFα (BioXCell clone XT3.11), 500μg anti-IFNAR1 (BioXCell clone MAR1-5A3) and/or 500μg IgG1 isotype control (BioXCell clone MOPC-21) in injection-grade saline. For inhibition of Nlrp3, mice were injected i.p. with 600μg MCC950 (SelleckChem #S7809) on the day of SCV2 infection and again two days later. For inhibition of arginase-1, mice were administered 100µg Nor-NOHA (Cayman Chemical #10006861) i.n. once daily on the day before, the day of, and two days following SCV2 infection.

### Viral quantification by TCID_50_ assay or RNA extraction and quantitative PCR of viral genomes

Viral quantitation was performed as previously described ^39,179^. Briefly, after harvesting lungs from mice, the inferior lobe, post-caval lobe and left lung were immediately homogenized in PBS for TCID_50_ assays. 10-fold serial dilutions were performed before plating on Vero E6 cells (American Type Culture Collection, #CRL-1586). TCID_50_ was calculated using the Reed– Muench method after 96 hours of incubation. To measure viral gene copy number, the superior lobe was homogenized in RLT Plus buffer (QIAGEN, #1053393) with β-mercaptoethanol following storage at −80°C in RNAlater (ThermoFisher, #AM7021). RNA was extracted from RLT Plus lysates using the RNeasy Plus Mini Kit (QIAGEN, #74136), including on-column DNase treatment using the RNase-Free DNase set (QIAGEN, #79256). The actively replicating (sub-genomic, sgRNA) conformation of the SCV2 E gene ^182^ was detected using primers at 500nM as follows: Forward (5’-CGATCTCTTG TAGATCTGTTCTC-3’), Reverse (5’-ATATTGCAGCAGTACGCACACA-3’) and the probe was used at 125nM (5′-(FAM)-ACACTAGCCATCCTTACTGCGCTTCG-(3IABkFQ)-3′). Copy numbers were calculated using a standard curve from a stock of known concentration ^137^.

### RNA sequencing and transcriptional analysis

RNA was extracted as described above and sent for sequencing by Novogene Corporation as previously described ^183^. Sequencing using the Illumina NovaSeq 6000 platform was performed to generate paired-end 150-bp reads. Sequences were aligned to the mouse transcriptome (GRCm38, mm10), comprising mRNA and ncRNA, using STAR ^184^ after quality control. The output from the mapping step was then converted to count tables using the tximport R package ^185^. The read count gene expression matrix was examined using the DESeq2 R package ^186^ to identify differentially expressed genes (DEG) in experimental groups. Changes in gene expression with a false discovery rate (FDR)-adjusted of p-value <0.05 and log2fold-change of ±1.3 were considered significant. Gene set enrichment analysis was then performed on the DEGs using the clusterProfiler R package ^187^ with the REACTOME database ^188^. DEGs common to both TLR-ligand pre-treated groups were entered into ImmGen’s MyGeneSet Browser (http://rstats.immgen.org/MyGeneSet_New/index.html) ^189^ to identify cell types in which those genes are commonly expressed based on the existing ImmGen ultra-low-input (ULI) cell-type specific RNA-Seq datasets. The entire gene expression dataset is available in Gene Expression Omnibus under accession no. GSE254993.

### Cell isolation for flow cytometry

Three minutes prior to euthanasia, mice were intravenously (i.v.) injected with 5 – 6μg per mouse of SuperBright 780 or BV711 labeled CD45 (30-F11) or CD45.2 (104) as previously reported ^64^. Lungs from infected mice were dissociated using scissors or a GentleMACS dissociator (Miltenyi Biotec) and cells were isolated and analyzed as previously described ^190^. The following antibody clones were purchased from Biolegend, Bio-Rad, R&D Systems, BD or ThermoFisher: anti-CD45.2 (clone 104), anti-CD45 (30-F11), anti-CD31 (390), anti-CD326 (G8.8), anti-CD24 (M1/69), anti-CD49f (GoH3), anti-I-Ab/I-E/MHC-II (M5/114.15.2), anti-podoplanin/PDPN/Gp38 (8.1.1), anti-Sca-1 (D7), anti-CD317/BST2/Tetherin (927), anti-Siglec-F (E50-2440 or 1RNM44N), anti-Ly6G (1A8), anti-CD68 (FA-11), anti-Ly6C (HK1.4), anti-CD11b (M1/70), anti-CD88 (20/70), anti-CD11c (N418 or HL3), anti-CD169 (SER-4), anti-TREM2 (237920), anti-CD64 (X54-5/7.1), anti-CD195/CCR5 (HM-CCR5, 7A4), anti-CD192/CCR2 (475301), anti-CD36 (HM36 or No. 72-1), anti-CD282/TLR2 (6C2), anti-arginase1/Arg1 (A1exF5), anti-Nos2 (CXNFT), anti-CD206 (C068C2), anti-CD38 (90), anti-Hif1α (241812), anti-LOX-1/OLR1 (214012), anti-ABCA1 (5A1-1422), anti-CD13 (R3-63), anti-CD14 (Sa14-2).

### Histopathology

The middle right lung lobe from each mouse was fixed in 4% paraformaldehyde, transferred to 70% ethanol, and paraffin-embedded before sectioning and mounting on glass slides for staining with hematoxylin and eosin (H&E). Stained slides were imaged by light microscopy on an Aperio Versa 200 (Leica). Images were processed using QuPath v0.3.2 and ImageJ v1.53t (NIH) as previously described ^39^.

### Multiplex cytokine array and ELISAs

Lung homogenates were prepared as described above for TCID_50_ assays and cytokines were measured using a ProcartaPlex Luminex kit (Thermo Fisher Scientific) on a MAGPIX Instrument (R&D Systems) according to the manufacturer’s instructions. Mouse and human ACE2 protein levels were quantified from lung homogenates by ELISA (R&D Systems #DY3437, #DY933).

### Statistical analyses

Statistical analyses were performed using Prism software version 9.0 for Mac OS X (GraphPad Software). The statistical details of experiments, including the statistical tests used, are listed within each figure legend. Outlier data points were identified and removed when n >10 using the ROUT method (Q=1%) in Prism. P values are indicated directly in the figures or are otherwise expressed as p < 0.05 (*), p < 0.01(**), p < 0.001 (***) with p values >0.05 considered not significant (n.s.). Data presented are combined of a minimum of two or more independent experiments unless otherwise stated in the figure legend.

## ACKNOWLEDGEMENTS

We are grateful to the dedicated staff of the Comparative Medicine Branch in the NIAID ABSL2 and ABSL3 facilities. We also thank Drs. A. Sher, I. Fraser, K. Fennelly, P. Murphy, C. Nelson, R. Namasivayam, M.Lionakis, A. Klion, E.V. Dang and S. Best for discussions and feedback on the manuscript. This work was supported by the Division of Intramural Research, National Institute of Allergy and Infectious Diseases (KDMB, DLB, RFJ). BBA and ATLQ are supported by the Intramural Research Program of the Oswaldo Cruz Foundation, Brazil, the National Institute of Allergy and Infectious Diseases [U01-AI069923], the Civilian Research and Development Foundation (CRDF) Global #DAA3-17-63144 and Brazilian National Council for Scientific and Technological Development (CNPq).

## AUTHOR CONTRIBUTIONS

Conceptualization: KDMB, PJB

Methodology: PJB, RFJ, KLH, JSK, KC

Investigation: PJB, ACB, EC, EPA, MSS, FTJ, STG, ATLQ, ERF, CMJ, KLH

Data analysis and visualization: KDMB, PJB, EPA, ATLQ, ERF, KLH,

Funding acquisition: KDMB, DLB, RFJ, BBA

Supervision: KDMB, DLB, RFJ, BBA

Resources: KDMB, DLB, RFJ, BBA

Writing - original draft: KDMB, PJB

Writing – review & editing: KDMB, PJB, DLB, ACB, EPA, RFJ, BBA, ATLQ, KLH

## SUPPLEMENTARY FIGURE LEGENDS

**Figure S1:**
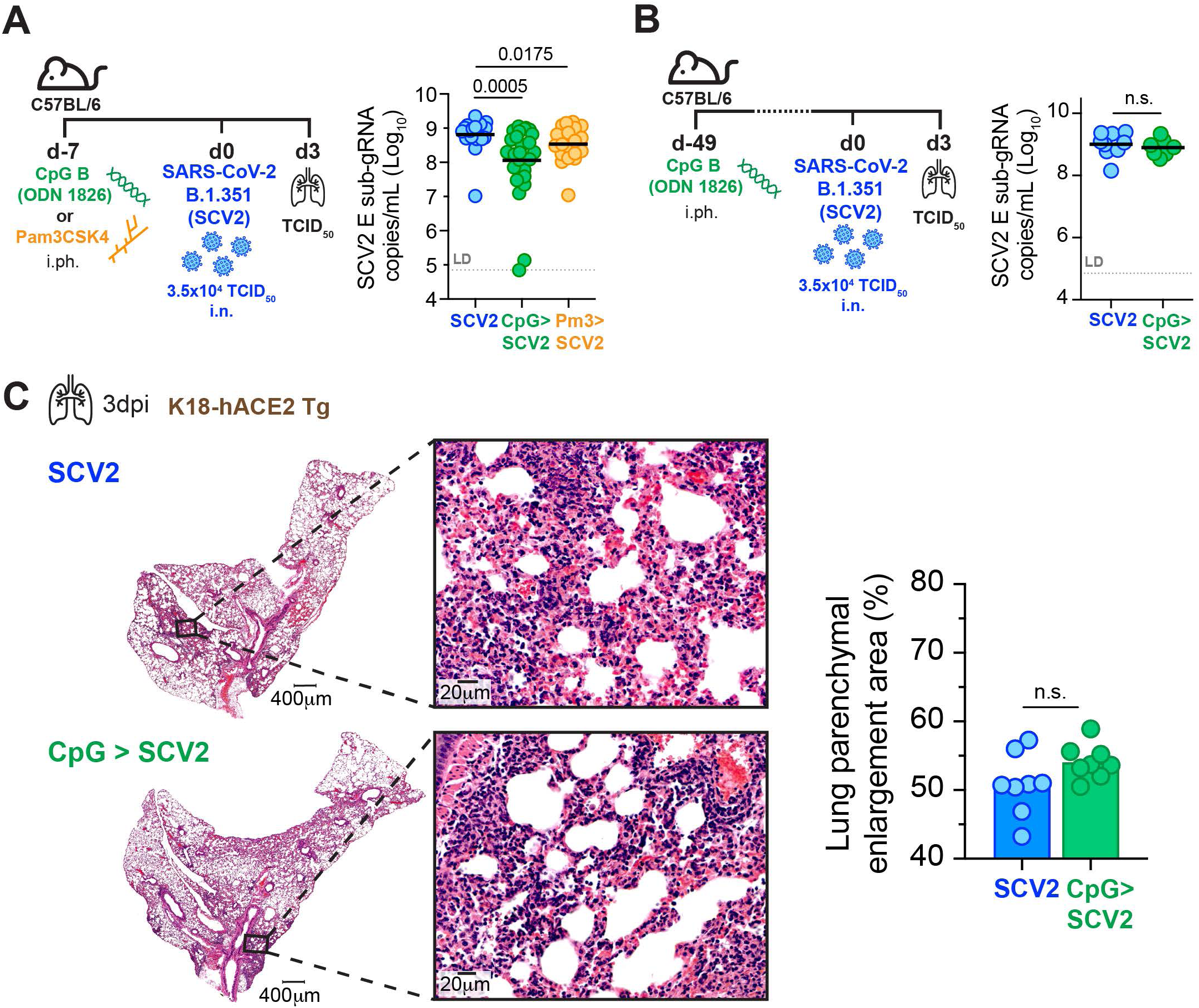
Recent one-time pulmonary TLR conditioning is sufficient to suppress SCV2 replication in the lung with no changes in gross lung pathology. **(A)** Schematic of WT mice given PBS, 10μg CpG or 50μg Pm3 intrapharyngeally (i.ph.) seven days prior to intranasal (i.n.) infection with 3.5×10^4^ TCID_50_ SCV2 (B.1.351) and SCV2 viral load in lungs as measured by qPCR for the SCV2 E gene in its sub-genomic form (sub-gRNA) at three days post-infection (3dpi), n= 19-25, data combined from five independent experiments. **(B)** Schematic of WT mice given either PBS or 10μg CpG i.ph. seven weeks before i.n. infection with 3.5×10^4^ TCID_50_ SCV2 (B.1.351), and SCV2 viral load in lungs as measured by qPCR for sub-gRNA SCV2 E gene on 3dpi, n= 9-10, data combined from two independent experiments. **(C)** Representative H&E staining of lung tissue from K18-hACE2 Tg mice given PBS or 10μg CpG i.ph. one week before infection i.n. with 1×10^3^ TCID_50_ SCV2 (USA-WA1/2020), mice were euthanized 3dpi (scale bars indicate magnification) and percentage of parenchymal enlargement was quantified, n= 8, data combined from two independent experiments. Geometric mean, significance determined by two-tailed Mann-Whitney test, LD= limit of detection, n.s.= not significant.

**Figure S2:**
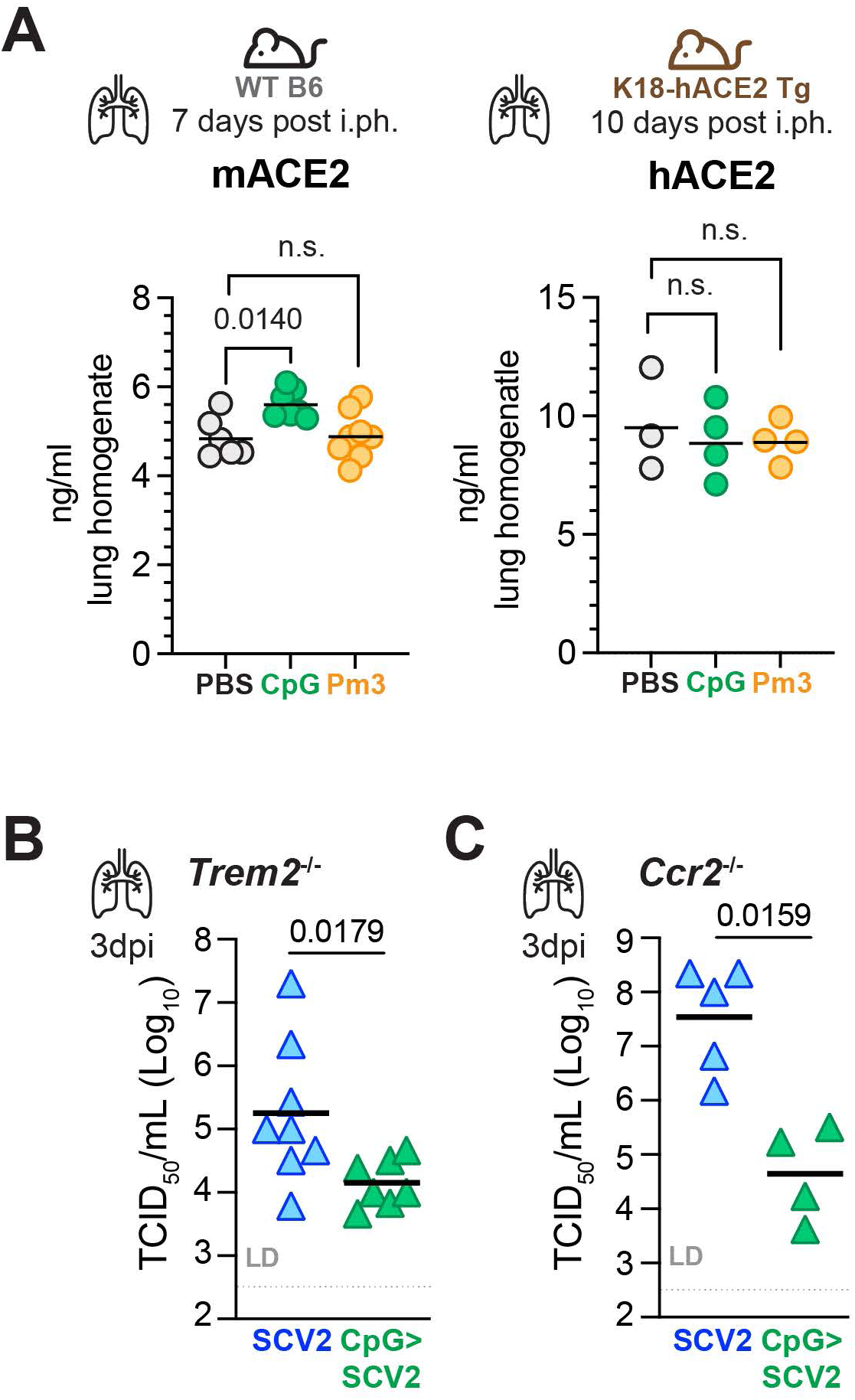
TLR-induced SCV2 restriction is not mediated through reduced ACE2 protein expression and is not reversed by deleting *Ccr2* or *Trem2*. **(A)** Left: WT mice were administered PBS, CpG or Pm3 intrapharyngeally (i.ph.). Lungs were collected at seven days post treatment and homogenates were assayed for mouse ACE2 by ELISA. Right: K18-hACE2 Tg mice were administered PBS, CpG or Pm3 i.ph., lungs were collected at 10 days post-treatment and homogenates were assayed for human ACE2 by ELISA, n= 3 – 8, data combined from 1 – 2 independent experiments. **(B)** *Trem2*^−/−^ or (C) *Ccr2*^−/−^ mice were given either PBS or 10μg CpG i.ph. seven days prior to being i.n. infected with 3.5×10^4^ TCID_50_ SCV2 (B.1.351), mice were euthanized 3 days later. Viral loads in lung are shown as measured by TCID_50_ on Vero E6 cells, n= 3 – 8, data combined from 1 – 2 independent experiments. Geometric mean, significance determined by two-tailed Mann Whitney test, LD= limit of detection, n.s.= not significant.

**Figure S3:**
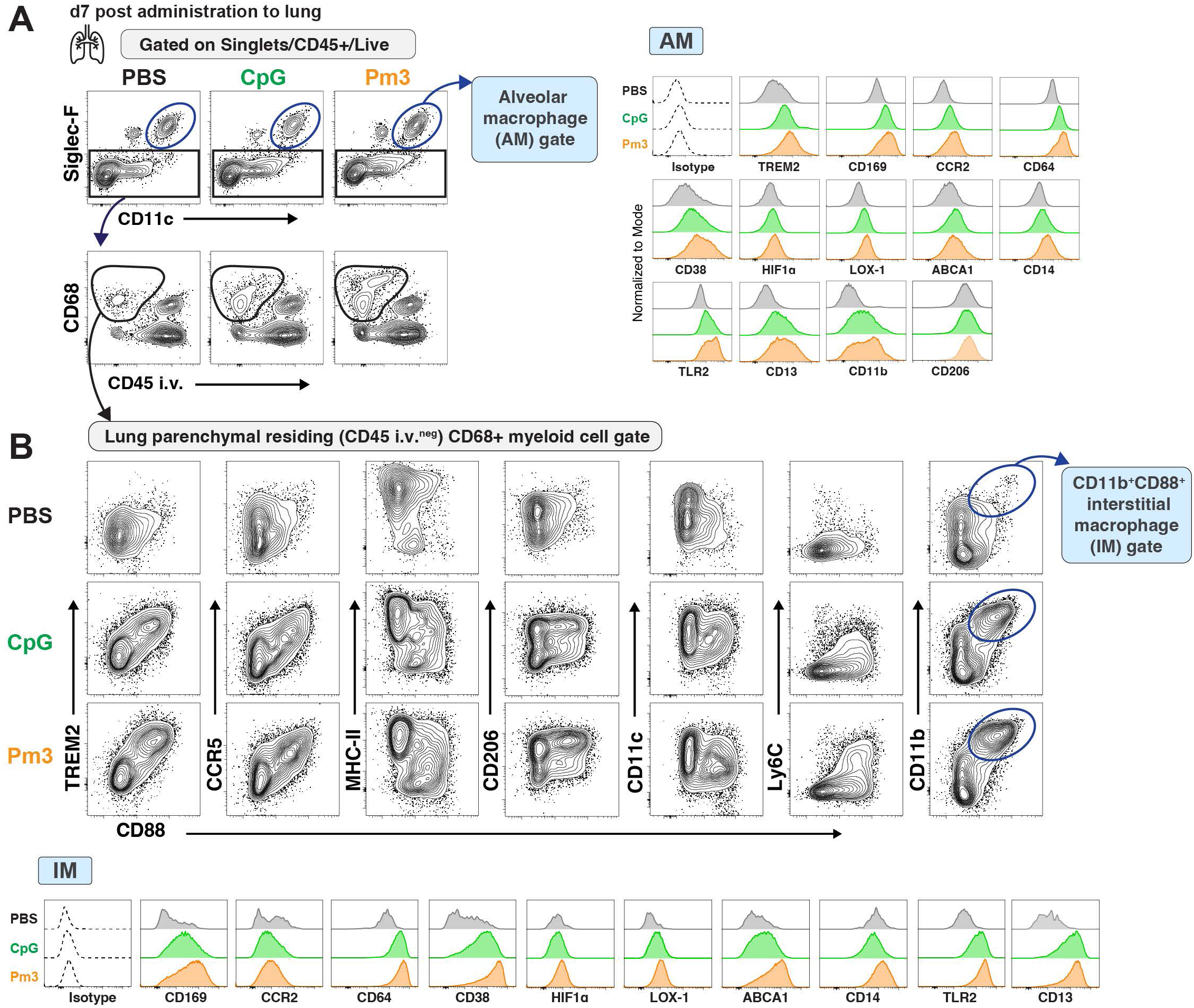
Recent pulmonary TLR pre-stimulation results in quantitative and qualitative changes to the tissue-resident macrophage compartment. **(A)** Example flow cytometry plots depicting gating strategy for identification of TRM related to **Fig 3**, and **Fig S4A** as alveolar macrophages (AM), and lung parenchymal (intravascular CD45 negative ( i.v.^neg^)) CD68^+^ interstitial macrophages (IM) cells from lungs of mice treated with 10µg CpG or 50µg Pm3 intrapharyngeally (i.ph.) seven days prior. Right: Histograms depicting relative expression of indicated markers on AM. **(B)** Example flow cytometry plots of CD11b^+^ CD88^+^ interstitial macrophages (IM) from the lung parenchymal residing CD68^+^ myeloid cell population identified in (A) showing the distribution of TREM2, CCR5, MHC-II, CD11c and Ly6C expressing cells within the lung parenchymal CD68^+^ myeloid cell population are also depicted. Histograms depicting the relative expression of indicated markers, n= 8 – 9, data combined from two independent experiments.

**Figure S4:**
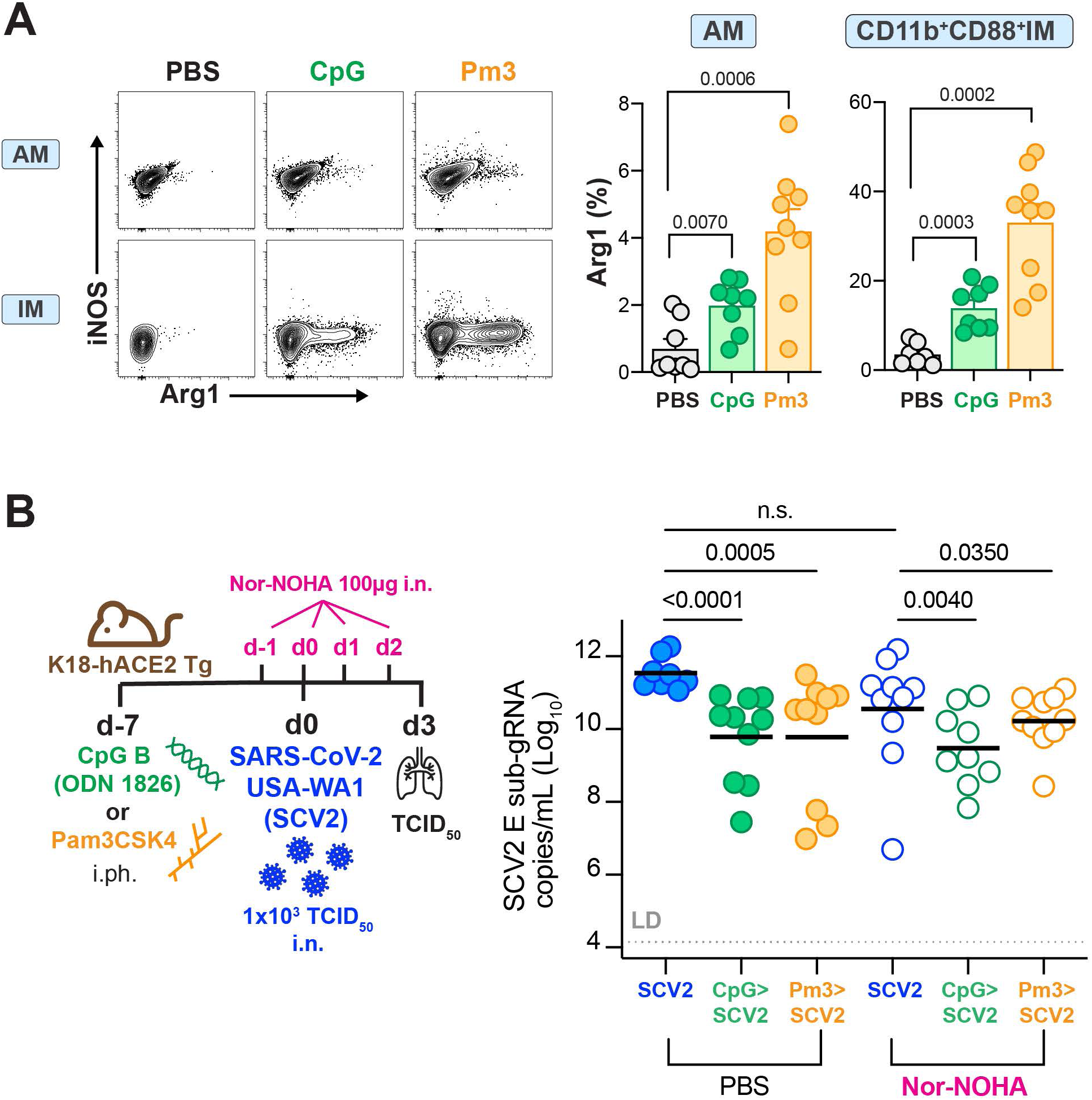
Arginase expression by TRM is elevated by recent pulmonary TLR stimulation but does not contribute to TLR-induced restriction of SCV2 viral replication. **(A)** Left: Example flow cytomwetry plots showing iNOS and arginase-1 (Arg1) expression from alveolar macrophages (AM, top) and CD11b^+^ CD88^+^ interstitial macrophages (IM, bottom) Right: Summary data of Arg1^+^ cells as a percentage of AMs (left panel) and IMs (right panel) n= 8 – 10, data combined from two independent experiments **(B)** Schematic of K18-hACE2 Tg mice given PBS, 10μg CpG or 50μg Pm3 intrapharyngeally (i.ph) sevemn days before intranasal (i.n.) infection with 1×10^3^ TCID_50_ SCV2 (SCV2, USA-WA1/2020) while given PBS or 100µg of the arginase inhibitor Nor-NOHA i.n. once daily from one day before SCV2 infection until two days after infection. Right: viral loads in the lung of PBS (filled circles) or Nor-NOHA (open circles) treated mice as measured by qPCR for the SCV2E gene in its sub-genomic form (sub-gRNA), n= 8 – 10, data combined from two independent experiments. Geometric mean, significance determined by two tailed Mann Whitney test, LD= limit of detection, n.s.= not significant.

**Figure S5:**
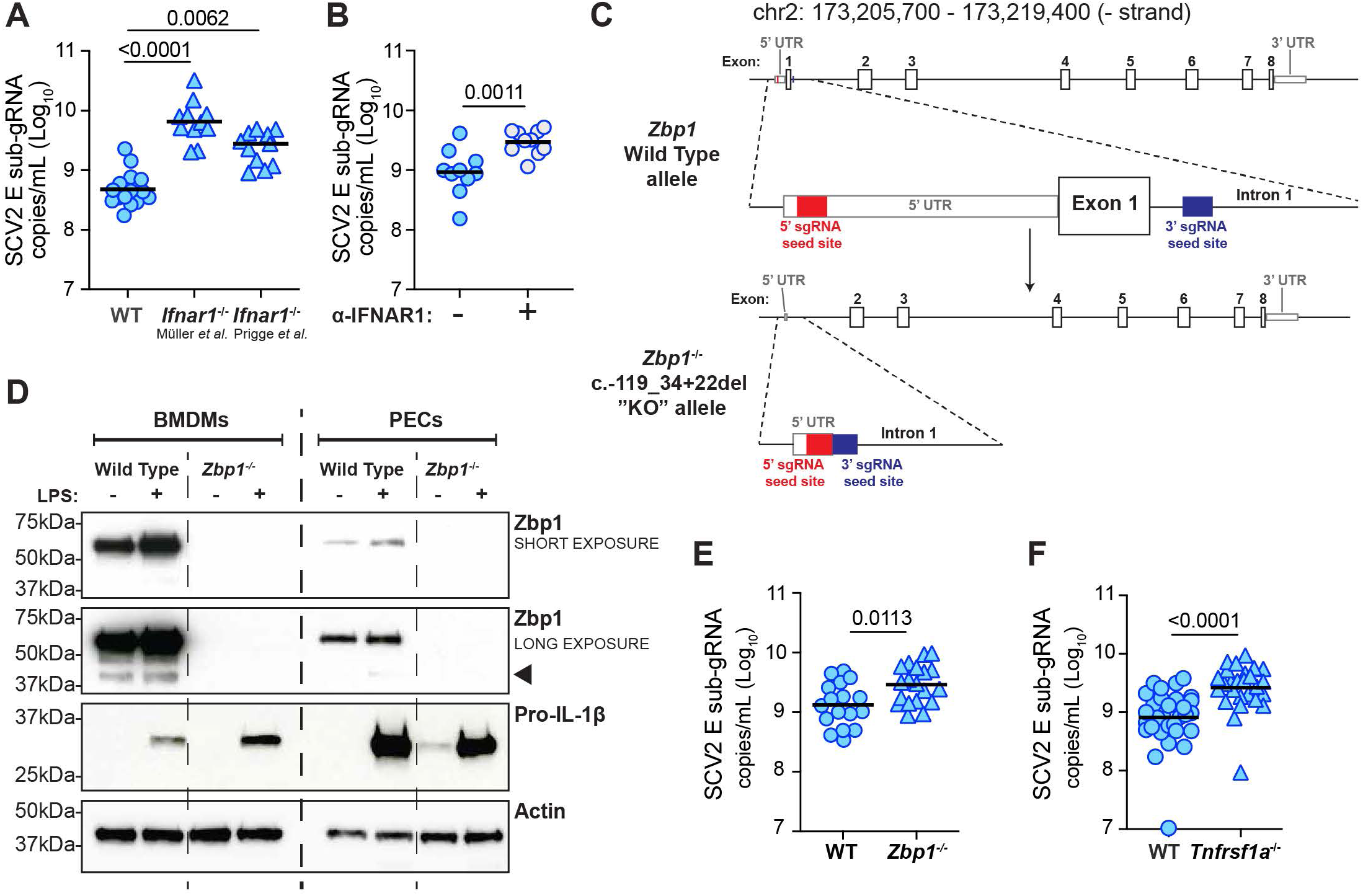
Description of the generation of *Zbp1^−/−^* mice and increased viral titers in *Ifnar1*, *Zbp1* and *Tnfrsf1a* deficient mice measured by qPCR for sub-genomic E genew. **(A – C)** Viral loads as measured by RT-qPCR for the SCV2 E gene in its actively replicating sub-genomic form (sub-gRNA) according to experimental setups shown in **Figure 4A** & **Fig 4B**: **(A)** two different strains of *Ifnar1*^−/−^ mice, **(B)** WT mice injected intraperitoneally with an anti-IFNAR1 monoclonal antibody. **(C)** Schematic showing the *Zbp1* gene (exons shown as white boxes with corresponding exon number), the binding sites for CRISPR sub-gRNAs used to create *Zbp1*^−/−^ mice (red and blue bars) and the resulting allele from deletion of the *Zbp1* 5’UTR and exon 1 by this strategy as confirmed by Sanger sequencing. **(D)** Western immunoblots of lysates prepared from bone marrow-derived macrophages (BMDMs) or peritoneal exudate cells (PECs) from either WT or the *Zbp1*^−/−^ mice. Cells were incubated with or without 200ng/mL LPS for six hours before collection for immunoblotting. Short and long exposures of Zbp1 are shown (the correct band for Zbp1 at 44kDa is indicated by the black arrow). Expression of pro-IL1β is included as a control for LPS stimulation and actin is shown as a loading control. **(E – F)** Viral loads as measured by qPCR for the SCV2 E gene in its sub-genomic (sub-gRNA) form in lung lysates from WT and **(E)** *Zbp1*^−/−^ or **(F)** *Tnfrsf1a*^−/−^ mice infected with SCV2 as described in **Fig 4A**, n= 9 – 29, viral titer data combined from 2 – 6 independent experiments, geometric mean, statistical significance calculated by two-tailed Mann Whitney test.

**Figure S6:**
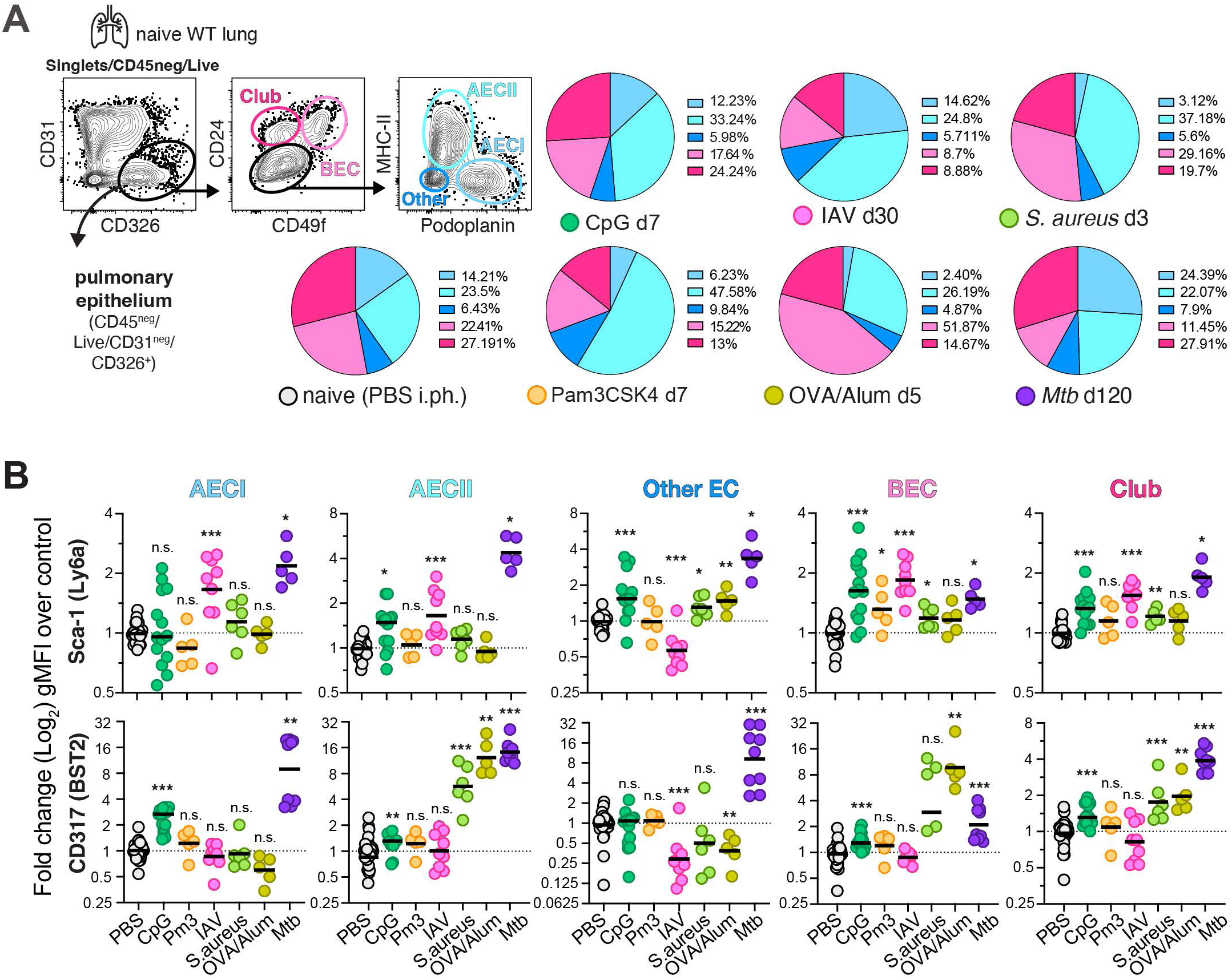
Prior pulmonary exposure to various inflammatory stimuli induces diverse remodeling of the lung epithelium. **(A)** Example flow cytometry plots from naive lungs of WT mice depicting the gating strategy for lung epithelial cell (EC) subsets and pie charts depict the proportion of epithelial cell subsets from of mice treated with various inflammatory or infectious stimuli compared to those from PBS control animals at the indicated time points without SCV2 (SCV2) infection, AEC (alveolar epithelial cells), BEC (bronchial epithelial cells), n=5-14, data is pooled from 2 – 4 independent experiments, for all conditions except OVA/Alum, which was done once **(B)** Fold change geometric mean fluorescence intensity (gMFI) of IFN-inducible surface marker (ISM) expression of Sca-1 and CD317 measured by flow cytometry on lung epithelial cell (EC) subsets from lungs of mice treated with various inflammatory or infectious stimuli compared to those from PBS control animals at the indicated time points without SCV2 infection, n=5-14, data is pooled from 2 – 4 independent experiments, for all conditions except OVA/Alum, which was done once. Geometric mean, significance calculated by two-tailed Mann Whitney test, * = p<0.05, ** = p<0.01, *** = p<0.001, n.s.= not significant.

